# TMEM63B functions as a mammalian thirst receptor

**DOI:** 10.1101/2024.02.01.578339

**Authors:** Wenjie Zou, Xingyu Chen, Jiamin Ruan, Siqi Deng, Huize Wang, Wuqiang Zhan, Jingxin Wang, Zhiyong Liu, Zhiqiang Yan

## Abstract

Thirst drives animals to reinstate water homeostasis by fluid intake. An increase of blood osmolality is thought to induce thirst by activating a thirst receptor expressed in the subfornical organ (SFO), but the molecular identity of this receptor remains elusive. Here, we provide behavioral and functional evidence to show that TMEM63B functions as a mammalian thirst receptor in the SFO and mediates osmotic and dehydrated thirst. First, we showed that TMEM63B is expressed in SFO excitatory neurons and required for the neuronal responses to hypertonic stimulation. Heterologously expressed TMEM63B is activated by hypertonic stimuli and point mutations can alter the reversal potential of the channel. More importantly, purified TMEM63B in liposomes establishes osmolarity-gated currents. Finally, Tmem63b knockout mice have profound deficits in thirst, and deleting TMEM63B within the SFO neurons recapitulated this phenotype. Taken together, these results provide a molecular basis for thirst 82and demonstrate TMEM63B is the long-sought mammalian thirst receptor.

## Introduction

Animals have the remarkable ability to sense their changing internal needs for water, and increased blood osmolality can trigger thirst^1–5^. Previously, the prevailing hypothesis was that thirst is primarily manifested as a sensation of dryness in the mouth and throat. However, our current comprehension demonstrates that thirst functions as a regulatory reaction to changes in the bloodstream. More precisely, the thirst is triggered when there is a rise in plasma osmolality or a reduction in plasma volume or pressure^6–10^. Thirst acts as a driving force for animals to search for and consume water, effectively bringing these blood metrics back to their normal physiological levels. The advent of novel tools for recording and manipulating neural activity has yielded profound insights into the neural circuits in the SFO, OVLT, and MnPO that underlie thirst and drinking behavior^11–19^. In contrast, the specific molecular components (thirst receptors) that convert blood osmolarity into neural signals still remain an enigma^4^. Changes in blood osmolality can be detected in the subfornical organ (SFO) and the organum vasculosum of the lamina terminalis (OVLT) in the forebrain^20–27^. These two sensory circumventricular organs (CVO) brain regions are located outside the blood-brain barrier and therefore have direct access to the circulation for plasma osmolarity sensing^27^. Specialized interoceptive neurons in the SFO and OVLT are intrinsically osmosensitive^28–31^. One hypothesis is that these SFO and OVLT interoceptive neurons monitor the blood osmolality directly through osmosensitive or mechanosensitive ion channels. Despite early studies suggesting the involvement of TRPV1 and TRPV4 cation channels in osmosensation^32–36^, subsequent work found that mice lacking the genes that encode these channels exhibit normal control of fluid balance and drinking behavior^4,37–40^. To date, the identity of the ion channels responsible for osmosensing in SFO and OVLT interoceptive neurons remains elusive^4^.

Excitatory neurons in the SFO are necessary and sufficient for the induction of thirst^11,13,14,18,21,22,27,41–43^, but it is still unclear whether SFO excitatory neurons are the interoceptive neurons that sense blood osmolarity changes directly. We first demonstrated that SFO excitatory neurons can be directly activated by hypertonic stimuli. Then, we examined the presence of recognized ion channels that respond to osmotic or mechanical stimuli, such as Piezo1/2, TMC1/2, and the TMEM63 family channel within these neurons. By analyzing the gene expression of SFO based on single-cell sequencing^17^, we revealed TMEM63B mRNA was expressed in SFO neurons. TMEM63B is a member of TMEM63/OSCA family of proteins that are conserved from plants to animals. Our previous study further showed that OSCA family proteins are mechanically activated ion channels^44^, and TMEM63 family proteins, the animal homologs of OSCA, were also shown to be mechano-gated ion channels^45^. The mammalian TMEM63 family has three members, TMEM63A, TMEM63B and TMEM63C, and their osmosensing ability and physiological function in mammals are worthy of studying^45–54^.

We then unveiled several significant discoveries about the roles of TMEM63B in thirst sensation. First, TMEM63B is prominently present in SFO excitatory neurons and plays a crucial role in their capability to perceive increases in osmolarity. Second, mice lacking TMEM63B and those with SFO-specific conditional knockout display a pronounced deficit in their ability to sense thirst. Of notable importance, when TMEM63B is expressed and isolated in a heterologous system, it forms an ion channel that detects hypertonic stimulation. These results show that TMEM63B functions as a mammalian thirst receptor, fulfilling the criteria for a sensory receptor has to meet^55,56^ and providing the molecular basis for behaviors associated with water homeostasis.

## Results

### SFO excitatory neurons can be directly activated by osmolarity changes and this activation requires TMEM63B

SFO excitatory neuron activation is necessary and sufficient for the generation of thirst^11,13,14,18,21,22,27,41–43^, but it is still unclear whether SFO excitatory neurons work as the primary sensory neurons to transduce osmolarity changes into neuronal signals and the sensory receptors in these neurons that mediate thirst also remain unknown^4^. To examine whether SFO excitatory neurons can be directly activated by hypertonic stimuli, physiological recordings of these neurons under hypertonic stimulation were carried out. In the osmotic and dehydrated model of thirst, the plasma osmolality in mice was approximately 350 mOsm/kg^17^. CaMK2α is a marker of excitatory neurons in the SFO^13^. We injected the rAAV2/9-CaMK2α-mCherry virus to label SFO excitatory neurons (Figure 1A). Action potentials were cell-attached recorded from SFO excitatory neurons labeled with red fluorescence under isotonic and hypertonic conditions (Figure 1B and Figure S1). SFO excitatory neurons show clear action potential firing to the hypertonic stimulus of 315 mOsm/kg and 350 mOsm/kg (Figure 1B and 1H). With hyperosmotic stimulation, we recorded the inward current in SFO excitatory neurons after blocking synaptic transmission (Figure 1C). Significant responses were recorded in SFO excitatory neurons under hypertonic stimulation of 315 mOsm/kg, and the higher hyperosmolarity of 350 mOsm/kg can elicit a current of approximately 30 pA (Figure 1J).

**Figure 1.**
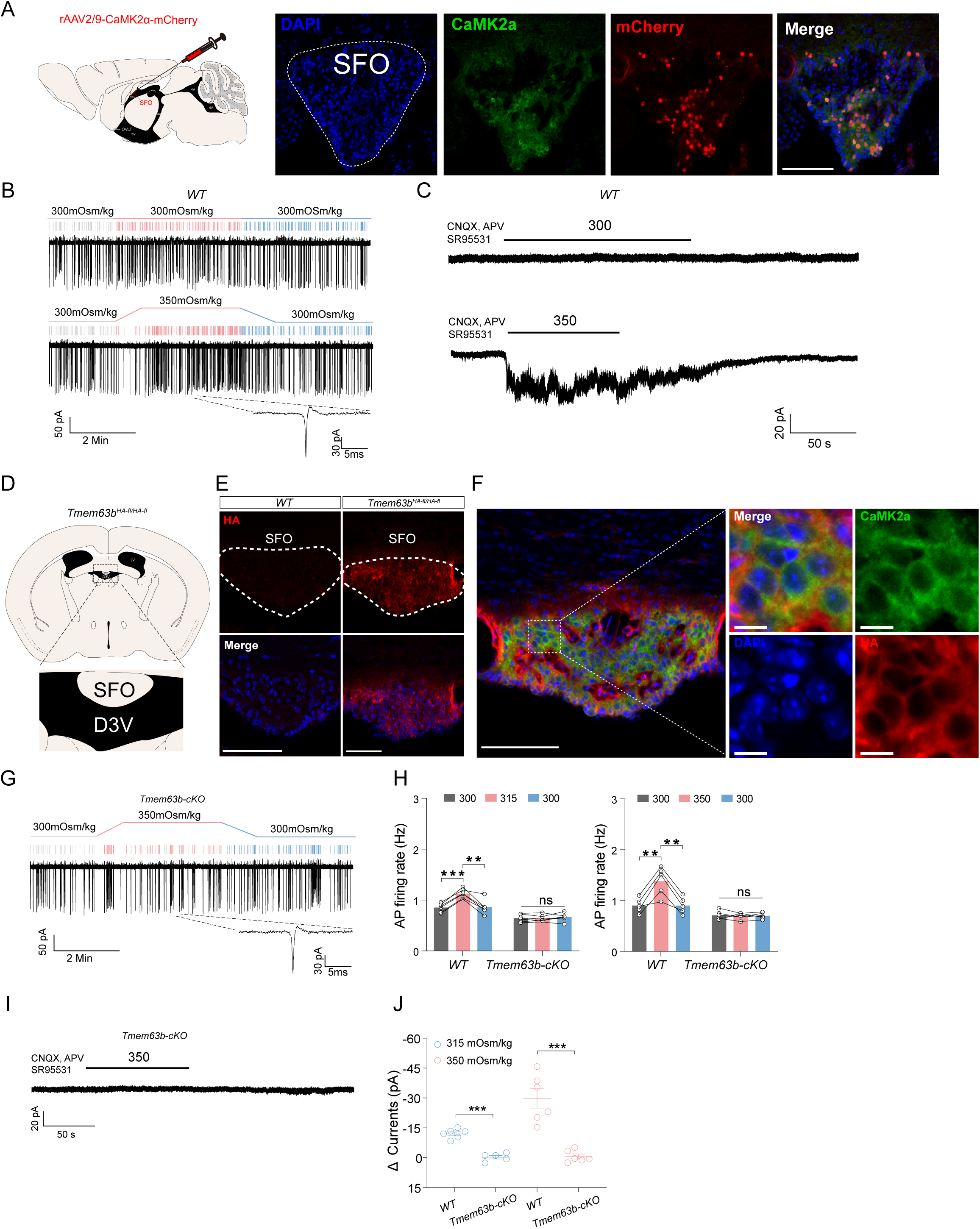
TMEM63B is required for the activation of SFO excitatory neurons in response to osmolarity changes. **(A)** Left, schematic diagram of labeling SFO excitatory neurons by injecting AAV-CaMK2α-mCherry in the SFO. Right, representative images show mCherry expression in SFO excitatory neurons. Scale bar 100 μm, mCherry (red) was expressed under the CaMK2α promoter. Green indicates endogenous CaMK2α, which labels excitatory neurons. **(B)** Representative traces of action potential (AP) firing of SFO excitatory neurons in wild-type (WT) mice. Isotonic (300 mOsm/kg) and hypertonic stimuli (350 mOsm/kg) were applied for 4 minutes (red). Raster potential firing of SFO excitatory neurons in response to isotonic or hypertonic stimuli. The firing rate of SFO excitatory neurons in WT mice shows a reversible increase in response to the hypertonic stimulus. **(C)** Representative traces of current recorded in SFO excitatory neurons in response to isotonic and hypertonic stimuli of 350 mOsm/kg. Significant currents were recorded 350 mOsm/kg. **(D)** The schematic diagram shows the anatomical location of the SFO (coronal view). D3V, dorsal third ventricle. **(E)** Endogenous TMEM63B is expressed in SFO. TMEM63B is labeled by a fusion HA tag in *Tmem63b^HA-fl/HA-fl^* mice. Red indicates immunofluorescence staining against HA, showing that TMEM63B is expressed in the SFO. Blue indicates nuclear staining. WT, wild type. Scale bar 100 μm. **(F)** TMEM63B labeled by a fusion HA tag (red) is highly expressed in CaMK2α-positive (green) excitatory neurons. Scale bar: left, 100 μm; right, 20 μm. **(G-H)** The AP firing of SFO excitatory neurons in response to hypertonic stimuli is dramatically reduced in *Tmem63b-cKO* mice. **(I-J)** Hyperosmosensitive currents induced by 315 mOsm/kg (left) or 350 mOsm/kg (right) were eliminated in *Tmem63b-cKO* mice. All error bars denote the mean ± SEM, \*\*\**P*<0.001, \*\**P*<0.01, *one-way* ANOVA and Student’s t test, for **H** and **J**, *n*= 5-7.

Based on the hypothesis that SFO excitatory neurons directly monitor blood osmolality by osmosensitive or mechanosensitive ion channels, we analyzed the expression of the known osmosensitive or mechanosensitive ion channels, including Piezo1/2, TMC1/2 and TMEM63 family channel in these neurons^45,48–50,53,54,57–68^, using published single-cell sequencing data collected from the SFO^17^. We found that TMEM63B mRNA is expressed in SFO excitatory neurons (Figure S2). To analyze TMEM63B protein expression in the SFO, we generated a polyclonal antibody against mouse TMEM63B, and found that TMEM63B is expressed in SFO (Figure S3A). To further validate the endogenous expression of TMEM63B in SFO, we used genetically edited *Tmem63b^HA-fl/HA-fl^* mice, in which the tandem hemagglutinin (HA) and 3×FLAG tags were inserted after the N-terminal signal sequence in exon 2 of the *Tmem63b* gene and two LoxP sites were inserted into intron 1 and intron 4 of the *Tmem63b* gene^48^. Immunofluorescence staining against HA or FLAG revealed TMEM63B is highly expressed in the SFO (Figure 1D, 1E and S3B), but not in OVLT and MnPO (Figure S4). Finally, by immunostaining for CaMK2α and HA in *Tmem63b^HA-fl/HA-fl^* mice, we found a majority of excitatory neurons (about 80%) in the SFO express TMEM63B (Figure 1F).

Next we injected the rAAV2/9-CaMK2α-Cre-mCherry virus into the SFO of *Tmem63B^HA-fl/HA-fl^* mice to selectively knock out TMEM63B in SFO excitatory neurons. Under hypertonic stimuli of both 315 and 350 mOsm/kg, the firing rate of SFO excitatory neurons was significantly increased in the control group, but this change was not observed in *Tmem63b-cKO* mice (Figure 1G and 1H). The hyperosmolarity-sensitive current was also lost in TMEM63B-deficient SFO excitatory neurons (Figure 1I and 1J), despite TMEM63B-deficient SFO excitatory neurons has normal response to glutamate (Figure S5A and S5B) and Ang II application (Figure S5C-S5F). These data indicate TMEM63B is required for osmotic sensing of excitatory neurons in SFO.

### Heterologously expressed TMEM63B in cultured cells responds to osmolarity changes

If TMEM63B is the primary sensor that mediates thirst in SFO excitatory neurons, it should be activated directly by hypertonic stimulation. To test this hypothesis, we examined whether heterologously expressed TMEM63B could form osmolarity-sensitive channels in culture cells. We first co-expressed human TMEM63B (HsTMEM63B) and GCaMP6f in N2a cells and monitored the calcium fluorescent intensity of transfected N2a cells during hypertonic stimulation. We stimulated the cells by changing the bath solution from an isotonic solution to a gradient of hyperosmotic solutions from 350 to 600 mOsm/kg. Compared with cells transfected with GCaMP6f alone (Figure S6), TMEM63B-expressing cells exhibited a significant calcium response under hyperosmolarity. The calcium fluorescence intensity increased as the hyperosmolarity increased (Figure 2A-C and 2E-I). The calcium signals of TMEM63B-expressing cells coincided with the osmolarity changes during solution perfusion (Figure S7). We also tested a hyposmolality of 220 mOsm/kg and found it induced a calcium influx at similar extend with that in 600 mOsm/kg hyperosmolarity (Figure 2D and 2I). These results show that heterologous expression of TMEM63B in N2a cells responds to a range of osmotic stimuli, indicating that TMEM63B has unique feature that enables it to respond to osmolality changes.

**Figure 2.**
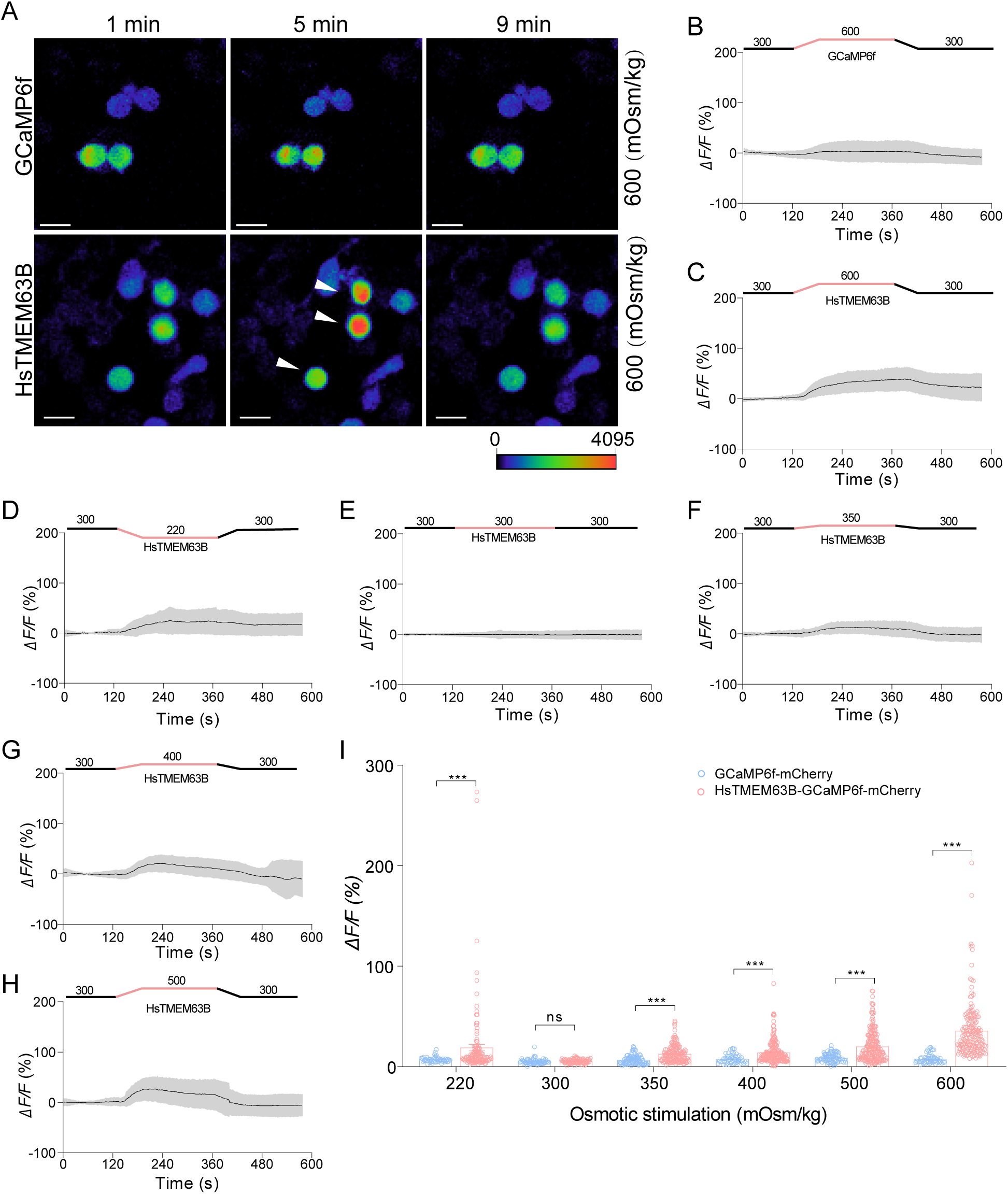
Heterologously expressed TMEM63B mediates calcium influx in response to hypertonic stimulus. **(A)** Representative images show calcium signaling responses in N2a cells under 600 mOsm/kg hyperosmotic stimuli, with heterologous expression of pCMV6-GCaMP6f- IRES-mCherry, or pCMV6-HsTMEM63B-P2A-GCaMP6f-IRES-mCherry. Arrowhead marks cells with cytosolic calcium increase. Scale bar 20 μm. **(B-C)** The calcium fluorescence intensity in response to the hypertonic solution (600 mOsm/kg). N2a cells heterologously were transfected with pCMV6-GCaMP6f-IRES-mCherry or pCMV6-HsTMEM63B-P2A-GCaMP6f-IRES-mCherry. The red line indicates the hyperosmotic stimulus for 4 minutes. **(D-H)** The calcium fluorescence intensity of N2a cells heterologously expressing pCMV6-TMEM63B-P2A-GCaMP6f-IRES- mCherry, in response to hyposmolarity of 220 mOsm/kg **(D)**, isotonic solution **(E)** and hyperosmolarity of 350 mOsm/kg **(F)**, 400 mOsm/kg **(G)**, 500 mOsm/kg **(H). (I)** Quantification of the calcium response in TMEM63B-expressing N2a cells in response to a range of osmotic stimulation, presented as an increase in fluorescence intensity after osmotic stimulation (*ΔF*) relative to that before stimulation (*F_0_*). As the degree of hyperosmolarity increased, TMEM63B-expressing N2a cells showed increased calcium signals. The gray shadow in the calcium imaging trace denotes the mean ± SD. All error bars denote the mean ± SD, n.s., nonsignificant difference, \*\*\**P*<0.001; Student’s t test.

To further validate that TMEM63B is an osmolarity-sensing ion channel, we recorded osmolarity-activated currents in CHO cells transfected with HsTMEM63B with hypertonic stimulus by patch clamp recording. The cells were recorded in isotonic solution (300 mOsm/kg) for 2 minutes, and then a gradient of hypertonic solution ranging from 320 mOsm/kg to 600 mOsm/kg was applied for 3 minutes before isotonic wash-out. To examine the hypertonic current responses at different voltage, a ramp protocol (-80 mV to +80 mV) was applied every two seconds. CHO cells transfected with HsTMEM63B show hyperosmolarity-induced currents (Figure 3A-3D) and the currents at -70 mV in the ramp protocol was plotted over time (Figure 3A and 3C). The magnitude of the hyperosmolarity-sensitive current in TMEM63B-overexpressing cells increased with the degree of stimulation (Figure 3G-3L). Hyperosmotic stimuli at 320 and 350 mOsm/kg close to the blood osmolality in osmotic or dehydrated animal models were able to induce osmosensitive currents in TMEM63B-expressing CHO cells (Figure 3H-3I, 3L); stronger hyperosmolarity of 400, 500 and 600 mOsm/kg induced a larger current in CHO cells expressing TMEM63B (Figure 3C, 3J-3L). In contrast, the control cells without TMEM63B transfection showed no hyperosmoticsensitive current (Figure S8). Additionally, the hyperosomotic-sensitive current under 500 mOsm/kg solution in TMEM63B-expressing CHO cells can be largely blocked by 100 µM ruthenium red (Figure 3E and 3F). These results suggest that TMEM63B can mediate osmolarity-induced current and is likely to be an osmolarity-sensing ion channel.

**Figure 3.**
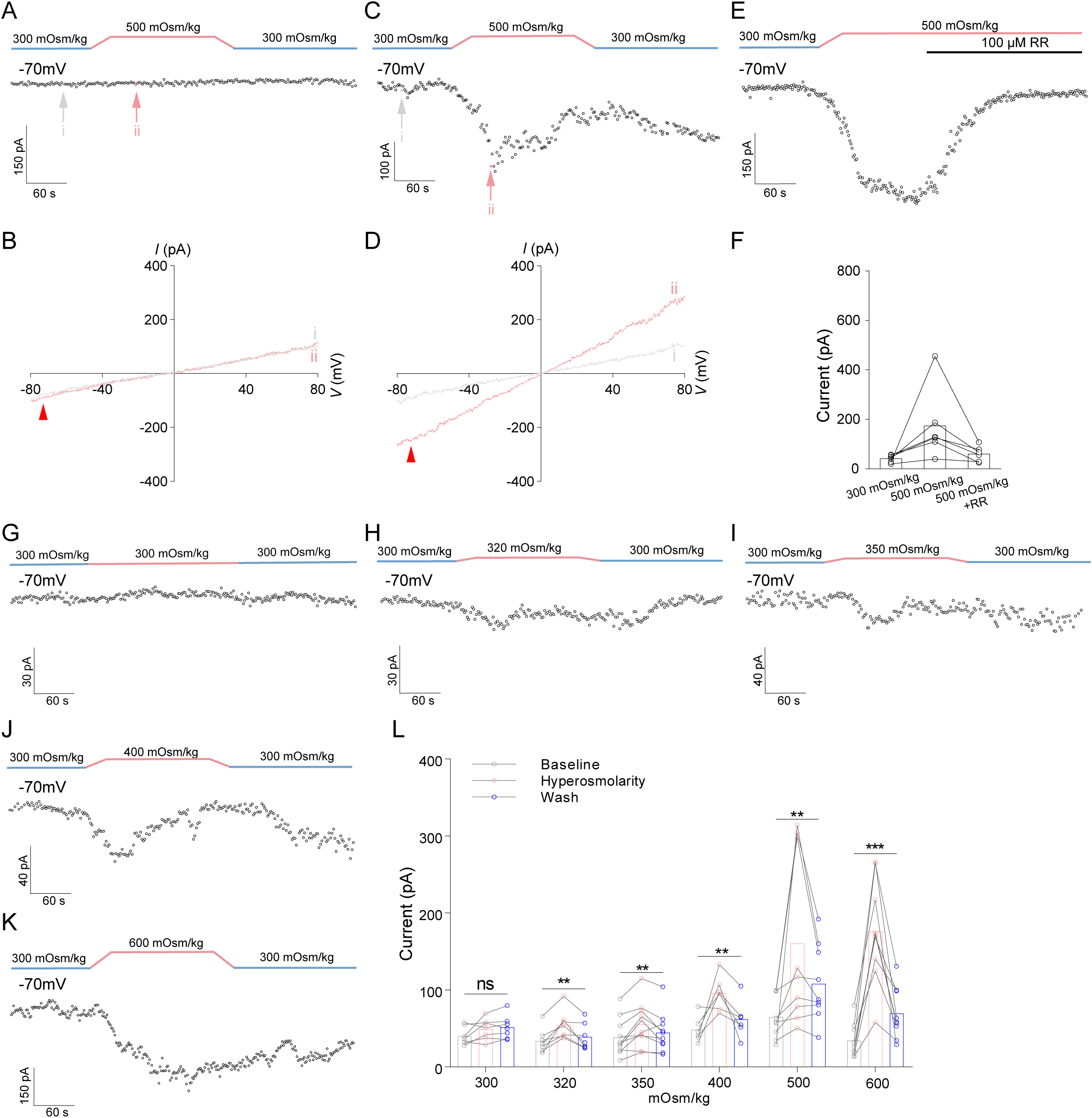
Heterologously expressed TMEM63B mediates hyperosmosensitive currents. **(A-B)** Representative whole-cell currents in untransfected CHO cells were recorded every two seconds by ramp protocols in 2-min isotonic solution (300 mOsm/kg; blue) and 3-min hyperosmotic solution (500 mOsm/kg; red) before isotonic wash-out. *I-V* curves at times indicated during hypertonic stress (red arrow in **A**) and isotonic solution (gray arrow in **A**) are shown in **B**. Currents at -70 mV (arrowhead in **B**) over time were plotted in **(A)**. **(C-D)** Representative whole-cell currents in CHO cells transfected with HsTMEM63B-eGFP were recorded in 2-min isotonic solution (300 mOsm/kg; blue) and 3-min hyperosmotic solution (500 mOsm/kg; red) before isotonic wash-out. *I-V* curves at times indicated during hypertonic stress (red arrow in C) and isotonic solution (gray arrow in **C**) are shown in **D**. Currents at -70 mV (arrowhead in **D**) over time were plotted in (**C**). **(E)** Representative whole-cell currents in CHO cells transfected with HsTMEM63B-eGFP were recorded in 2-min isotonic solution (300 mOsm/kg; blue) and 3-min hyperosmotic solution (500 mOsm/kg; red) with the application of ruthenium red (RR). Currents at -70 mV over time were plotted. **(F)** The current induced by hyperosmolarity in TMEM63B-expressing CHO cells can be largely blocked by RR. **(G-K)** Representative whole-cell currents in CHO cells transfected with HsTMEM63B-eGFP in response to isotonic solution **(G)** and hyperosmolarity of 320 mOsm/kg **(H)**, 350 mOsm/kg **(I)**, 400 mOsm/kg **(J)**, 600 mOsm/kg **(K). (L)** The maximum hyperosmolarity-induced currents at -70 mV in CHO cells expressing HsTMEM63B-eGFP increased with the gradient of evaluated hyperosmotic stimuli (320, 350, 400, 500 and 600 mOsm/kg). n.s., nonsignificant difference, \*\**P*<0.01, \*\*\**P*<0.001; *one-way* ANOVA; n = 5-10.

### TMEM63B is likely a pore-forming subunit of the hyperosmosentive channel

Although N2a and CHO cells heterogeneously expressing HsTMEM63B exhibit strong sensitivity to hyperosmotic stimulus, whether TMEM63B is a pore-forming subunit of osmosensitive channel remains unknown. To answer this question, we firstly built two homologous structures of HsTMEM63B by SWISS-MODEL and AlphaFold and mapped a putative pore region in the predicted structures (Figure 4A), based on previously published structures of the plant OSCA/TMEM63 channels. By analyzing the structure and alignment of the multiple homologous members of TMEM63B (Figure S9), we found a conserved glutamic acid residue (E510) in the putative pore region, which is negatively charged and may play a role in selectivity of the cation selective ion channel TMEM63B (Figure 4B). Then, we generated alanine and lysine substitution at this, E510A and E510K, respectively, and expressed them in CHO cells to record hyperosmolarity-induced currents. To analyze the effects of these two mutations, we recorded the whole-cell currents and switched the bath solution from an isotonic solution (300 mOsm/kg) to a hyperosmotic solution (500 mOsm/kg). The currents at -70mV were plotted to show the electrophysiological response of TMEM63B-transfected cells to the osmolarity change (Figure 4C and 4D). E510A showed a clear response to hypertonic stimulus, while the current response was barely detectable in the E510K mutant (Figure 4C and 4D). E510A also shifted reversal potential of the current induced by a hypertonic stimulus (Figure 4D and 4E), suggesting that TMEM63B is likely a pore-forming subunit of an osmolarity-sensing channel.

**Figure 4.**
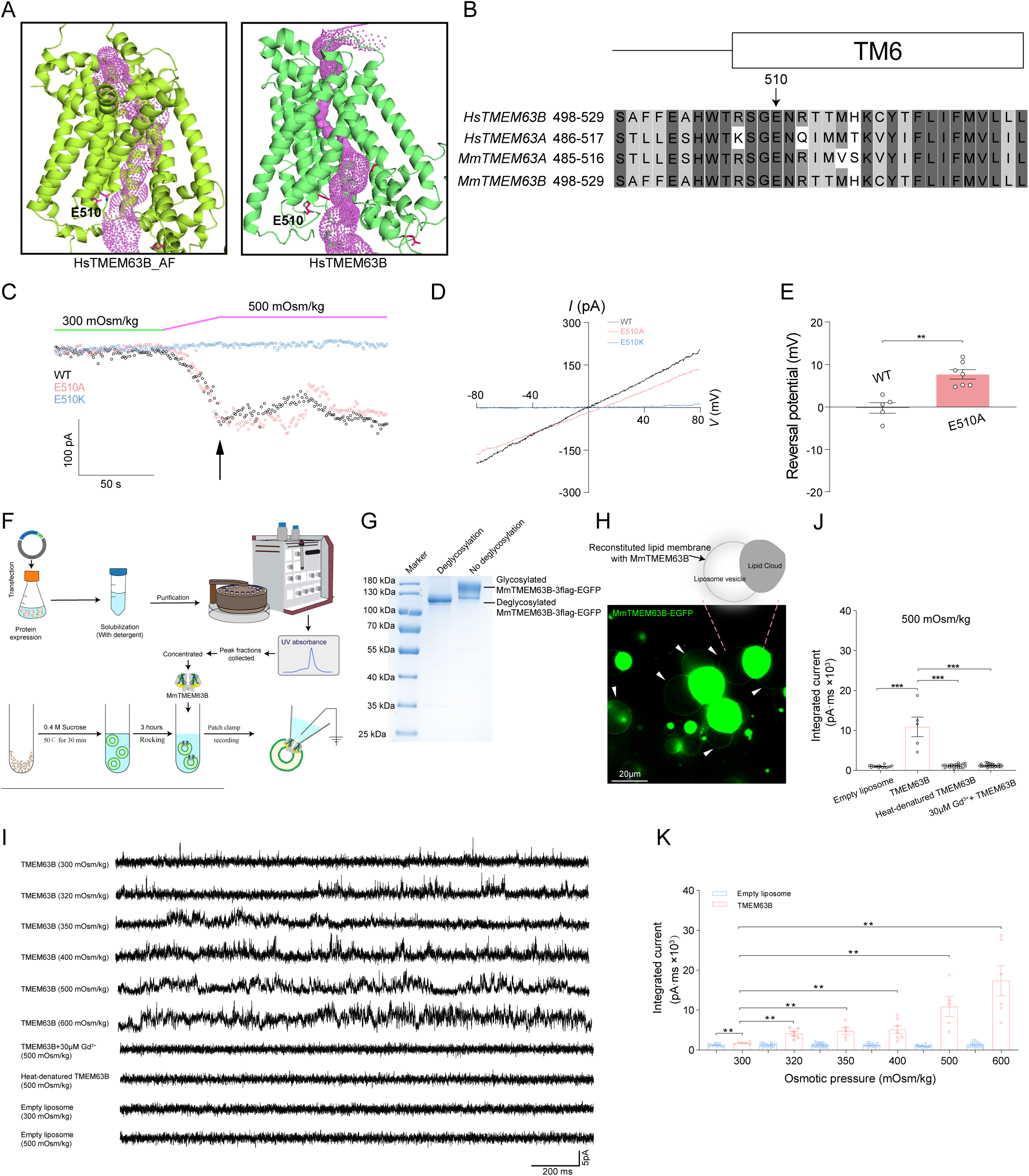
TMEM63B is likely a pore-forming subunit of a hyperosmosentive channel. **(A)** A side view of the human TMEM63B transmembrane domain was obtained from an AlphaFold model (left) or a homology model based on OSCA 1.2 (PDB: 6ijz, right). The calculated pore profile of the putative HsTMEM63B is indicated as purple dots. **(B)** Sequence alignment of human TMEM63B and its homologs. The arrow marks the negatively charged E510 residues in the putative pore region targeted for mutagenesis. Mm, *Mus musculus*; Hs, *Homo sapiens*. **(C)** Currents at -70 mV over time induced by a 3-min 500 mOsm/kg hyperosmolarity stimulus were plotted for representative CHO cells transfected with wild-type or the two putative selectivity filter mutants (E510A and E510K) of HsTMEM63B. Whole-cell currents were monitored every two seconds by ramp protocols from -80 mV to 80 mV for 100 ms. **(D)** The E510A mutation altered the *I-V* curve of hyperosmolarity-induced currents, whereas the E510K mutation largely eliminated the osmosensitive current under hypertonic stimulus. **(E)** The E510A mutation shifted the reversal potential of TMEM63B currents activated by a hypertonic stimulus. For **(C-E)**, error bars denote the mean ± SEM, \*\**P*<0.01, \*\*\**P*<0.001, Student’s unpaired *t* test, *n* = 5-7. **(F)** Schematic diagram of size-exclusion chromatography (SEC) for MmTMEM63B protein purification and proteoliposome reconstitution procedure with the sucrose method for patch-clamp recording. **(G)** SDS‒PAGE analysis of purified MmTMEM63B proteins with an eGFP tag stained with Coomassie blue. The deglycosylation treatment resulted in a significant reduction in the band smearing observed in SDS‒PAGE. **(H)** Representative images show eGFP-labeled MmTMEM63B incorporated into liposome membranes (bottom) and a carton illustration (top). The white arrows indicate membranes with eGFP-labeled MmTMEM63B. Scale bar, 20 μm. **(I)** Representative currents of MmTMEM63B incorporated into liposome membranes in response to isotonic or a gradient of hypertonic solutions. No current was recorded in the empty liposomes, TMEM63B-reconstituted liposomes with 30 µM GdCl_3_ or proteoliposomes of heat-denatured TMEM63B in 500 mOsm/kg hyperosmotic solution. All recordings were performed under +40 mV. **(J)** Quantification of integrated current responses in empty liposomes, MmTMEM63B-reconstituted proteoliposomes with or without 30 µM GdCl_3_ and the proteoliposome of heat-denatured TMEM63B in 500 mOsm/kg hyperosmotic solution holding at +40 mV. The area under the curve indicates the integrated currents. **(K)** Quantification of integrated current responses in empty liposomes and MmTMEM63B-reconstituted proteoliposomes under isotonic or a gradient of hypertonic conditions holding at +40 mV. \*\**P*<0.01, \*\*\**P*<0.001, Student’s t test. All error bars denote the mean ± SEM. The response to 500 mOsm/kg of empty liposomes and TMEM63B reconstituted liposomes and in **(J)** and **(K)** were same group of data.

To test whether purified TMEM63B proteins were sufficient to recapitulate the channel properties recorded from CHO cells transfected with TMEM63B, we reconstituted the purified MmTMEM63B proteins into liposomes (Figure 4F). The purified MmTMEM63B-3flag-eGFP (127.8kDa) migrated as smeared bands on SDS-PAGE (Figure 4G). Our previous study shows that the *At*OSCA1.1 protein, a homologous protein of TMEM63B in plants, was glycosylated when expressed in HEK293F cells^44^. To test whether mouse TMEM63B was also glycosylated when expressed in HEK293F cells, we treated the purified MmTMEM63B-3flag-eGFP with a protein deglycosylation mixture, the treatment notably diminished the smear in the bands (Figure 4G). Mass spectrometry examination of the purified MmTMEM63B-eGFP identified three N-glycosylated amino acid sites (Figure S10). These results indicated that the smeared bands in SDS-PAGE were due to glycosylation rather than other proteins. Next, we incorporated MmTMEM63B into liposomes for patch clamp recordings^63^, and eGFP-tagged MmTMEM63B localizes in the liposome membrane (Figure 4H).

The currents were recorded in the patches excised from empty liposomes and liposomes reconstituted with MmTMEM63B in the isotonic or hypertonic application of from 320 to 600 mOsm/kg. We found that the duration and current amplitude in MmTMEM63B reconstituted liposomes were significantly increased in the hypertonic stimulus (Figure 4I and 4K), with a gradient of elevated hyperosmolarity ranging from 320 to 600 mOsm/kg, an increasing integrated current of TMEM63B in liposomes was recorded (Figure 4I and 4K). The hyperosmolarity-induced currents in the TMEM63B-reconstituted liposomes under 500 mOsm/Kg were eliminated by 30 μM GdCl_3_ (Figure 4I and 4J). As further negative controls, empty liposomes or liposomes constituted with heat-denatured TMEM63B also failed to produce channel activity under hyperosmolarity stimulus (Figure 4I and 4J). These results suggested that TMEM63B is likely a pore-forming subunit of the hyperosmosensitive channel.

### TMEM63B is required for the drinking behavior of mice in the osmotic thirst and dehydrated thirst model

Having found that TMEM63B is an osmolarity sensitive channel and identified the functional role of TMEM63B in excitatory neurons activation by hypertonic stimuli in brain slices, we tested whether TMEM63B is required for the activation of SFO excitatory neurons in thirst models *in vivo*. After osmotic thirst or dehydrated thirst modeling (Figure 5A), control and *Tmem63b* knockout mice were sacrificed for immunostaining using c-Fos as a marker of thirst-evoked neuronal activity within the SFO. Consistent with previous studies^13,17^, we found SFO excitatory neurons are robustly activated, as indicated by c-Fos expression under osmotic thirst or dehydrated thirst, and this activation was greatly reduced in *Tmem63b* knockout mice (Figure 5B-5G). These results indicate that TMEM63B is required for the SFO excitatory neuron activation in thirsty condition.

**Figure 5.**
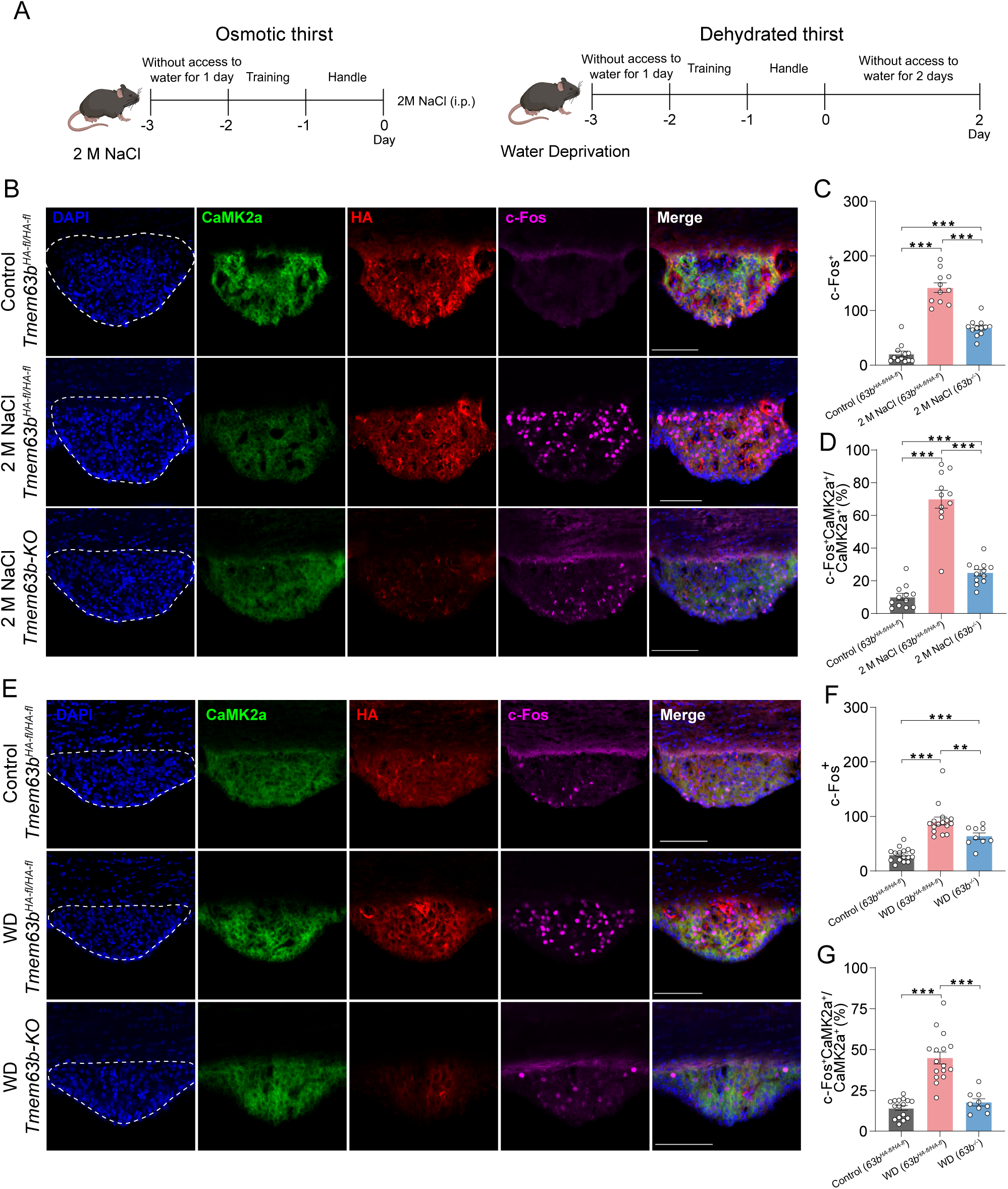
c-Fos immunostaining showed that TMEM63B is required for the activation of SFO excitatory neurons by osmotic or dehydration stress. **(A)** Schematic diagram and experimental procedures of the osmotic or dehydrated thirst model. i.p., intraperitoneal injection. **(B) and (E)** Representative images of the SFO region from *Tmem63b^HA-fl/HA-fl^* and *Tmem63b* knockout mice stained with antibodies against HA (red), CaMK2α (green), and c-Fos (purple) under osmotic **(B)** and dehydrated thirst **(E)** conditions. WD, water deprivation. Scale bar 100 μm. **(C) and (F)** Quantification of c-Fos-positive SFO neurons under both osmotic thirst **(C)** and dehydrated thirst **(F)** conditions. The number of c-Fos-positive cells increased under osmotic or dehydrated thirst conditions in *Tmem63b^HA-fl/HA-fl^* mice, and these increases were greatly reduced in *Tmem63b* knockout mice. **(D) and (G)**, The co-labeling of c-Fos and CaMK2α in c-Fos-positive excitatory neurons. The percentage of c-Fos-positive excitatory neurons increased dramatically under osmotic **(D)** or dehydrated thirst **(G)** conditions in *Tmem63b^HA-fl/HA-fl^* mice, and these increases were greatly reduced in *Tmem63b* knockout mice. For (**B-G)**, Control (*63b^HA-fl/HA-fl^*) represents the untreated *Tmem63b^HA-fl/HA-fl^* mice. 2 M NaCl (*63b ^HA-fl/HA-fl^*) or WD (*63b ^HA-fl/HA-fl^*) denotes the *Tmem63b^HA-fl/HA-fl^* mice under osmotic or dehydrated thirst conditions. 2 M NaCl (*63b^-/-^*) or WD (*63b^-/-^*) indicates *Tmem63b^-/-^* mice under osmotic or dehydrated thirst conditions. All error bars denote the mean ± SEM, \*\**P*<0.01, \*\*\**P*<0.001, *one-way* ANOVA, for (**C-F**), *n*= 9-16.

Having found that TMEM63B is required for the activation of SFO excitatory neurons during thirst stress and that TMEM63B is a pore-forming subunit to detect osmolarity changes, to finally determine whether TMEM63B is the mammalian thirst receptor, we investigated if TMEM63B plays a role in mediating thirst behavior in mice. We examined the drinking behavior of *Tmem63b* knockout mice in the osmotic thirst model and dehydrated thirst model, employing a two-choice consumption preference assay^1,11,17,41^. For the osmotic thirst model, mice were injected with 2 M NaCl intraperitoneally. For the water deprivation thirst model, mice were deprived of water for 2 days. For the two-choice consumption preference assay, mice in the osmotic thirst model or dehydrated thirst model were tested for the choice of water or 0.3 M NaCl (Figure 6A). Consistent with previous studies^1,11,17,41^, acute osmotic stress with hypertonic solution induced selective consumption of water over the 0.3 M NaCl solution (Figure S11A). Moreover, the number of licks and the drinking volume of water of *Tmem63b* knockout mice were greatly reduced compared with those of wild-type mice (Figure 6B). A similar phenotype of *Tmem63b* knockout mice was found in the water deprivation model (Figure 6C and S11B), indicating that TMEM63B is required for both osmotic and dehydrated thirst.

**Figure 6.**
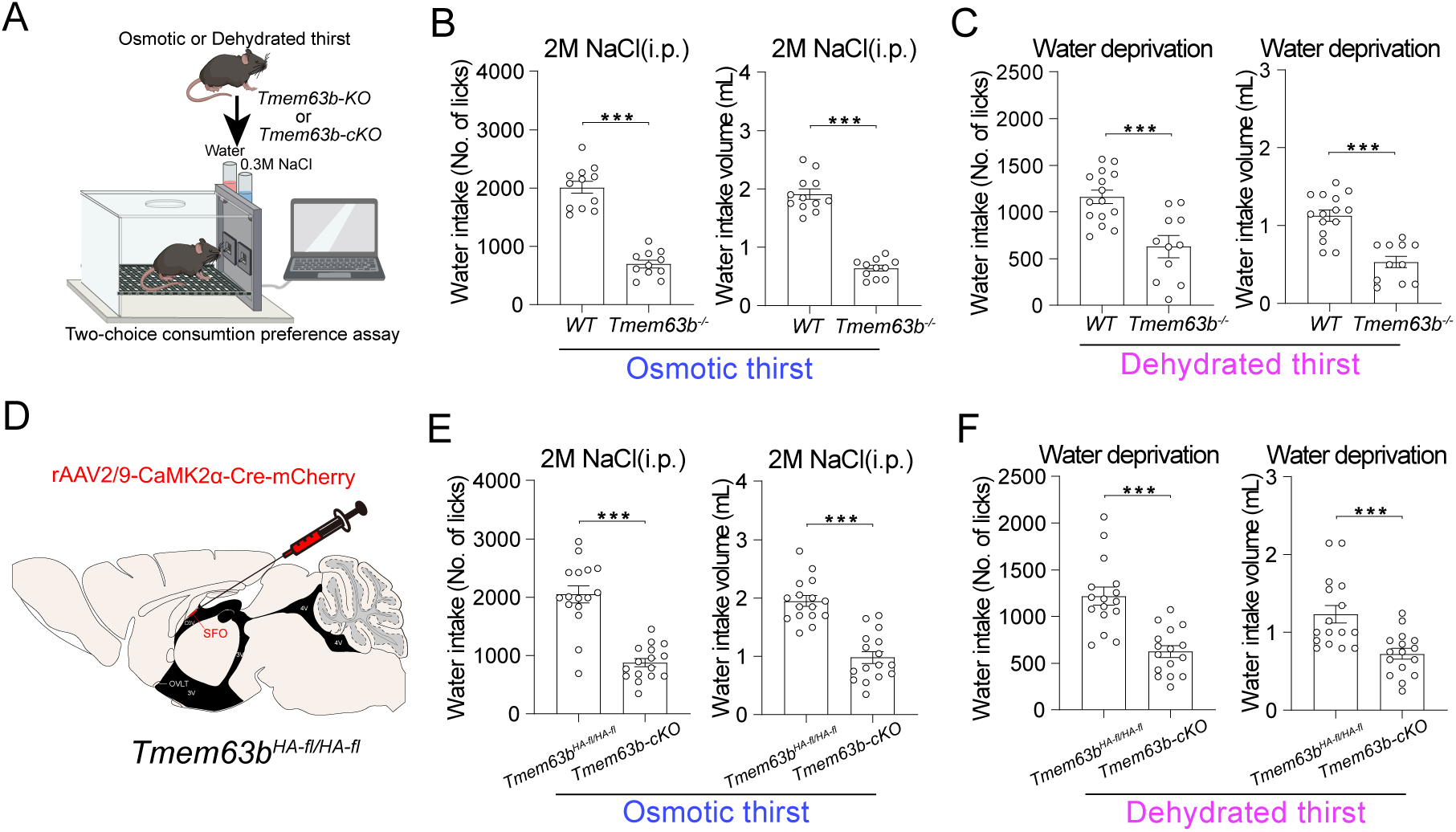
Conditional deletion of TMEM63B in SFO excitatory neurons profoundly impairs thirst behavior in mice. **(A)** Schematic diagram and experimental procedures of a two-choice consumption preference assay of osmotic or dehydrated thirst model mice. i.p., intraperitoneal injection. **(B)** Under osmotic thirst conditions, water intake of wild-type or *Tmem63b* knockout (*Tmem63b^-/-^*) mice during a 1-h session. The number of licks and intake volume of water in *Tmem63b* knockout mice were significantly reduced. **(C)** The water intake of wild-type or *Tmem63b^-/-^* dehydrated model mice during a 1-h session is shown. **(D)** Schematic diagram of inactivating TMEM63B in SFO excitatory neurons by injecting AAV-CaMK2α-Cre-mCherry in the SFO to generate *Tmem63b-cKO* mice. **(E)** The water intake of *Tmem63b^HA-fl/HA-fl^* or *Tmem63b-cKO* osmotic thirst model mice during a 1-h session is shown. The number of licks and intake volume of water in *Tmem63b-cKO* mice were significantly reduced. **(F)** The water intake of *Tmem63b^HA-fl/HA-fl^* or *Tmem63b-cKO* dehydrated thirst model mice during a 1-h session is shown. All error bars denote the mean ± SEM, \*\**P*<0.01, \*\*\**P*<0.001, Student’s unpaired *t* test, *n*=11-16.

To determine the role of TMEM63B in SFO excitatory neurons in thirst behavior, we injected the rAAV2/9-CaMK2α-Cre-mCherry virus into the SFO of *Tmem63B^HA-fl/HA-fl^* mice to selectively knock out TMEM63B in SFO excitatory neurons (Figure 6D). Behavior tests were conducted after thirst modeling. In both the osmotic thirst model and dehydrated thirst model, *Tmem63b-cKO* mice consumed less water than WT mice (Figure 6E and 6F, Figure S11C and S11D), demonstrating that TMEM63B deficiency in SFO excitatory neurons impairs thirst in mice. These results indicate that TMEM63B in SFO excitatory neurons is required in thirst.

## Discussion

Most mammals maintain a body fluid osmolality with a narrow range of approximately 300 mOsm/kg^73^. Under the osmotic or water deprivation model of thirst, the plasma osmolality in mice is approximately 350 mOsm/kg^17^. When body fluid osmolality increases, the animals feel thirsty and instinctively seek and consume water, restoring their body fluid osmotic pressure and plasma volume within the homeostatic range. The advent of novel techniques for neural recording and manipulation has yielded remarkable discoveries concerning the neural circuits involved in thirst and drinking behavior^11–19,74–77^. However, in contrast, the molecular identities of the primary thirst receptors responsible for converting changes in blood osmolarity into neural signals remain unknown^4^.

For a channel to be considered a bona fide transducer of blood osmolality to thirst (thirst receptor), it should meet several criteria^55^. TMEM63B satisfies all of these criteria. First, it is expressed in SFO excitatory neurons that are critical for thirst and TMEM63B is required for the drinking behavior of mice in thirst model; second, it is required for hypertonic solution-induced excitation of SFO excitatory neurons; third, heterologous expression of TMEM63B in cultured cells generates osmolarity-sensitive channels; fourth, purified TMEM63B shows hypertonic-induced currents. These results demonstrated that TMEM63B is a thirst receptor in mammals. Thirst is critical for water homeostasis, and conceivably, a deficiency of TMEM63B signaling might result in deficit of water homeostasis and lead to hypertension or kidney diseases.

## Acknowledgments

We thank Yun Stone Shi at Guangdong Institute of Intelligence Science and Technology for the *Tmem63b^HA-fl/HA-fl^* mice. We thank Yulong Li at Peking University, Boxun Lu at Fudan University and Zhou-Feng Chen in Shenzhen Bay Laboratory for comments. We thank the Bio-imaging Core of Shenzhen Bay Laboratory for providing imaging support. We also thank the Laboratory Animal Core of Shenzhen Bay Laboratory for providing mouse breeding and maintenance support.

## Author Contributions

W.Z. performed patch-clamp recording in cultured cells and brain slices, behavior tests and calcium imaging. X.C. mainly performed immunofluorescence staining, behavior tests, calcium imaging, J.R. purified MmTMEM63B, conducted patch-clamp recording in recombinant proteoliposomes, S.D. performed patch-clamp recordings experiments in cultured cells, H.W. performed calcium imaging, Wuqiang Zhan purified MmTMEM63B, J.W. and Z.L. performed single cell RNA-seq data analysis, Z.Y. initiated and supervised the study, Z.Y., W.Z. and designed the experiments, Z.Y., S.D. and W. Z. wrote the manuscript.

## STAR Methods

### KEY RESOURCES TABLE

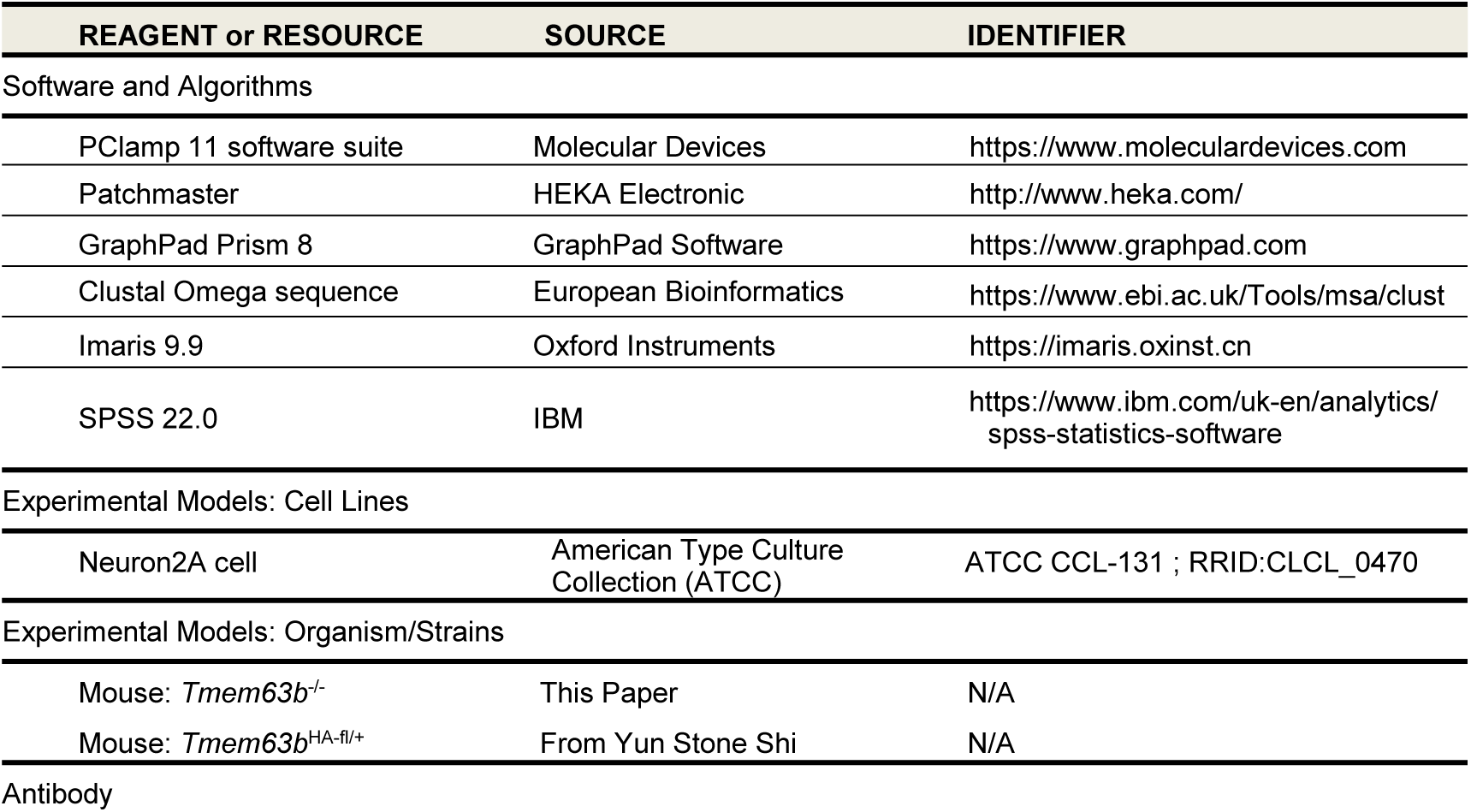
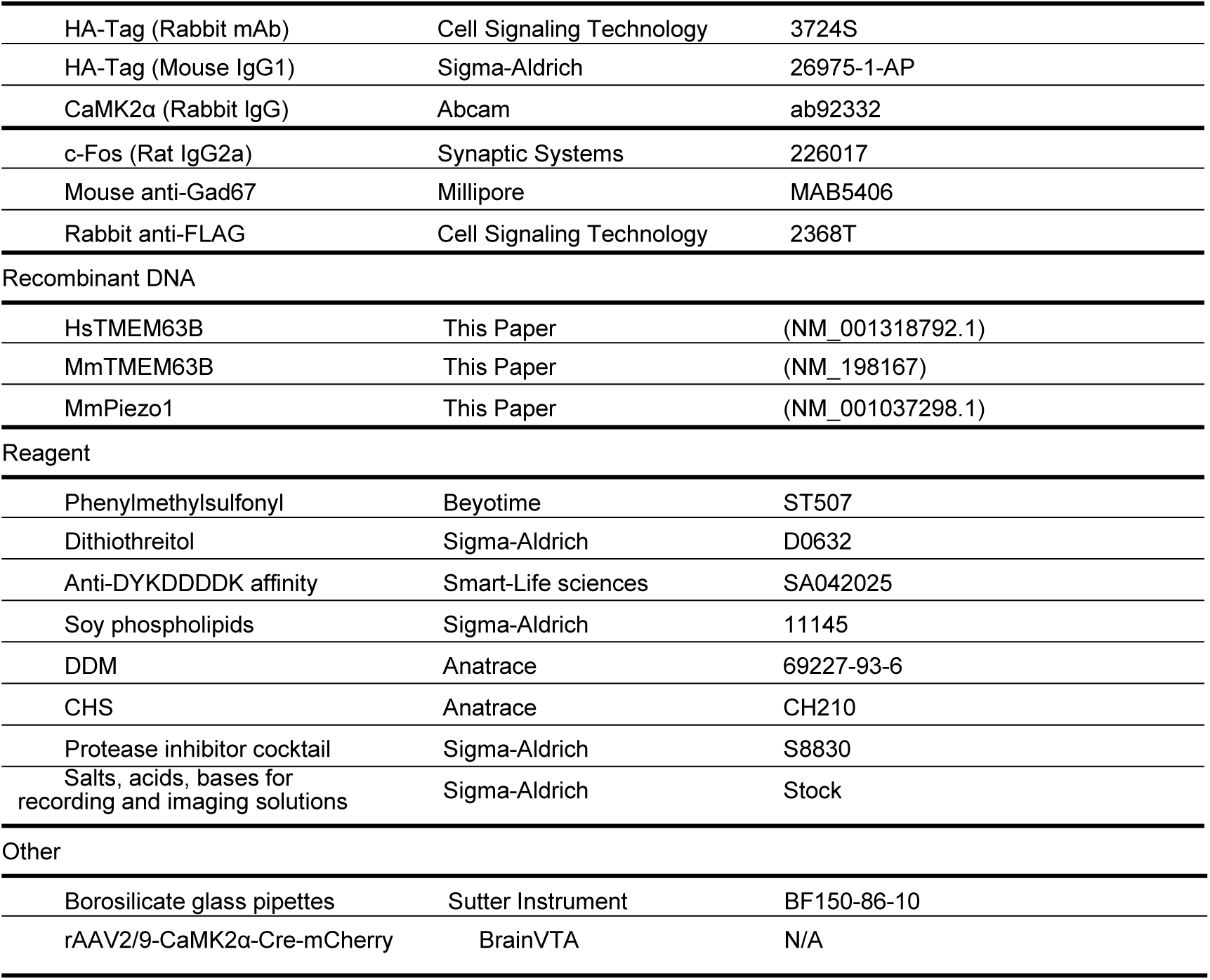

## Resource Availability

### Lead Contact

Further information and requests for resources and reagents should be directed to and will be fulfilled by the Lead Contact, Zhiqiang Yan.

### Materials Availability

All plasmids and genetically modified mice are available upon request.

### Data and Code Availability

The raw data presented in the study are available from the Lead Contact upon request.

## Method Details

### Animals

Mice were bred and maintained in standard cages under a condition of 12-hours light-dark cycle in a specific-pathogen-free (SPF) animal facility accredited by the Association for Assessment and Accreditation of Laboratory Animal Care (AAALAC). All the animal experiments involving the operation on mice were rigidly conducted by the animal protocols, approved by the Institutional Animal Care and Use Committee of the animal facility of the Model Animal Research Center of Shenzhen Bay Laboratory, Shenzhen, China. Both male and female animals were unbiasedly used in all the animal experiments. Mice from 8-12 weeks old were used in this study. Detailed information is indicated in the figures and legends. *Tmem63b^HA-fl/+^* mice were given by Yun Stone Shi^48^. *Tmem63b* knockout (*Tmem63b^-/-^*) mice were generated by mating *Tmem63b^HA-fl/+^* and CMV-Cre (Jackson NO. 006054) mice.

### Cell culture

Neuro2a (N2a) cells were grown in Dulbecco’s Modified Eagle Medium (DMEM) containing 4.5 mg/ml glucose, and 10% fetal bovine serum (GIBCO, USA). CHO cells were grown in DMEM/F12 containing 15 mM HEPES and 10% fetal bovine serum. Cells were plated onto 8-mm square glass poly-L-lysine coated cell slides placed in 35 mm dishes and transfected using Lipofectamine 3000 (Invitrogen) according to the manufacturer’s instructions. All plasmids were transfected at a constant concentration of 1000 ng/ml. The full-length coding sequence of human TMEM63B (NM_001318792) was amplified from HEK293 cell cDNA. The mouse *Tmem63b* gene (NM_198167) was synthesized by GenScript.

### Induction of physiological states and behavioral assays

For the osmotic thirst experiments, the mice were water deprivation for 24 hours to train in the Skinner box and were handled the next day. Then, the mice were injected with 2 M NaCl (5 μl/g body weight) and subjected to 10 minutes of water and food deprivation before behavioral testing. For the c-Fos staining, mice were kept in the home cage without access to food or water for 1 h to allow for stimulus-induced expression of immediate early genes.

For the water deprivation experiments, the trained and handled mice were fed food and 1 ml of water daily for 48 hours. After that, the mice were subjected to behavioral testing or euthanized for c-Fos staining.

All of the behavioral assays were performed in a custom Skinner box (Beijing Zhongshi Dichuang Technology Development Co. , Ltd) with different paradigms unless otherwise noted. The mice were trained in the Skinner box under water deprivation for 24 hours before the behavioral assays to train. As previously described^17^, for the two-choice consumption preference assay, one bottle of water and one bottle of 0.3M NaCl were presented in sequence during the same session. Mice in different groups were recorded for 1 hour, and the number of licks toward different solutions was measured.

### Ca^2+^ imaging in N2a cells

Cytoplasmic calcium levels were monitored using GCaMP6f. To form a tandem expression system, GCaMP6f was added to the C-terminus of HsTMEM63B via a P2A linker (TMEM63B-P2A-GCaMP6f-IRES-mCherry). The pCMV6-TMEM63B-P2A-GCaMP6f-IRES-mCherry vector was transfected into Neuro-2a cells prepared on the cell slide, and the pCMV6-GCaMP6f-IRES-mCherry vector was used as a control. After a 48 h transfection, the cells were perfused with extracellular solutions of varying osmolarities at a constant exchange speed using a peristaltic pump. Calcium fluorescence images were obtained at 0.5 Hz for 10 min using a laser scanning confocal microscope (Nikon A1, Japan) with a 20X objective lens and an excitation wavelength of 488 nm and 561 nm at room temperature (24 ± 2 °C). Cells were recorded in the isotonic solution for 2 min, switched to hypertonic solution for 4 min, and then switched back to isotonic solution for another 4 min. The isotonic solution (300 mOsm/kg) contained (mM): 72 NaCl, 5 KCl, 1 CaCl_2_, 1 MgCl_2_, 10 HEPES, 130 mannitol, pH 7.4 adjusted with NaOH. Additional hyperosmotic solutions were prepared by adjusting the solutions with mannitol without changing the ion concentrations. The osmolarity of the solutions was measured using a freezing point osmometer (Gonotec, 3000 Basci, Germany). Time-lapse images of the N2a cells were analyzed to acquire the fluorescence intensity in the region of interest (ROI) in each frame. The change in fluorescence intensity *ΔF/F_0_* was calculated as *100%*(F_t_-F_0_)/F_0_*, where *F_t_* was the fluorescence intensity at time t and *F_0_* was the average fluorescence intensity prior to the application of the osmolarity stimulus.

### Immunohistochemistry

Mice were deeply anesthetized with pentobarbital sodium (50 mg/kg) and perfused with PBS followed by 4% paraformaldehyde (PFA, pH 7.4). Mouse brains were extracted, then fixed overnight at 4 °C in 4% PFA and coronally sectioned at 35 μm intervals using a freezing microtome (CM3050S, Leica). Brain sections were permeabilized with a solution containing 5% bovine serum albumin and 1% Triton X-100 in PBS for 40 minutes at room temperature. Then the sections were blocked with a solution of 5% bovine serum albumin in PBS for 2 hours at room temperature followed by overnight primary antibody (1% bovine serum albumin in PBS) at 4 °C. The following primary antibodies were used: rabbit anti-HA-Tag (1:100, Cell Signaling Technology 3724S), mouse anti-HA-Tag (1:500, rabbit Sigma-Aldrich H3663), anti-FLAG (1:50, CST 2368T), rabbit anti-CaMK2α (1:1000, Abcam ab92332), mouse anti-Gad67 (1:500, Millipore MAB5406), rat anti-c-Fos (1:2500, Synaptic Systems 226017), and rabbit anti-TMEM63B (1:1000, custom-made affinity-purified polyclonal antibody with antigen sequence of TDADRLRRQERERVC). After washing three times with PBS, the brain sections were stained with secondary antibody for 2 hours at room temperature. The following secondary antibodies (1:1000, Thermo Fisher Scientific) were used: Alexa Fluor 488 Goat anti-Mouse (A11029), Alexa Fluor 488 Goat anti-Rabbit (A11008), Alexa Fluor 568 Goat anti-Mouse (A11031), Alexa Fluor 568 Goat anti-Rabbit (A11036), Alexa Fluor 647 Goat anti-Rat (A48265). DAPI (5 μg/ml) was then incubated for 5 min at room temperature. Following three additional washes with PBS, the brain sections were mounted on glass slides and imaged on a confocal microscope (Nikon A1). Cell counting was performed using Imaris (version 9.9).

### TMEM63B-eGFP expression and purification

The *MmTmem63b* (NM_198167) gene was synthesized and subcloned and inserted into the vector pCDNA3.1-eGFP. A 3 × FLAG tag (DYKDHDGDYKDHDI DYKDDDDK) was added to the C-terminus of the *MmTmem63b* gene with a GSAGS linker. The MmTMEM63B protein used for reconstitution was expressed in HEK293F cells. For protein expression and purification, HEK293F cells were grown in SMM 293-TII medium to a density of 1.8×10^6^ cells/mL and transfected with 1 μg DNA and 3 μl 1 mg/mL PEI in 20 mL of Opti-MEM. HEK293F cells were cultured for 60 hours at 37℃ and then harvested and pelleted. The pellets were washed with ice-cold PBS and stored at -80 °C for future use. The pellets were thawed on ice and sonicated in 50 mM Tris-HCl (pH 7.5), 150 mM NaCl with 1 mM phenylmethylsulfonyl fluoride, protease inhibitor cocktail (Sigma, S8830) and 1 mM dithiothreitol (DTT, Sigma D0632). Then, the cell lysate was incubated with SuperNuclease at 4 °C for 5 minutes, 2% DDM and 0.4% CHS were added, and the solutions were stirred for 2 hours. Insoluble material was removed by ultracentrifugation for 45 min at 40000 g (Beckman, JA-25.50 rotor), and the supernatant was loaded onto Anti-DYKDDDDK affinity beads (Smart-Life Sciences, SA042025). After the beads were washed with 50 mM Tris (pH 7.5), 150 mM NaCl, 0.05% DDM, 0.01% CHS, and 1 mM DTT, the protein was eluted with 50 mM Tris (pH 7.5), 150 mM NaCl, 0.05% DDM, 0.01% CHS, 200 μg/mL 3 × FLAG peptide, and 1 mM DTT. The sample was concentrated using a 100 kDa MWCO Amicon Ultracentrifugal filter (Millipore, UFC910096) and then applied to a Superose^TM^ 6 Increase 10/300 GL (Cytiva, 29091596) equilibrated with a buffer containing 50 mM Tris-HCl (pH 7.5), 150 mM NaCl, 0.05% DDM, 0.01% CHS, and 1 mM DTT. The peak fractions corresponding to MmTMEM63B-eGFP were concentrated at 1 mg/mL. The proteins were aliquoted and stored at -80 °C.

### Reconstitution of MmTMEM63B-eGFP in liposomes

Purified proteins were incorporated into artificial liposomes, which were formed as previously described^63,78^. A 200 μL 20 mg/mL solution of soy phospholipids (Sigma, 11145) in chloroform was dried in a glass tube under a stream of N_2_ while rotating to form a homogeneous lipid film. Once dry, 5 μL of ddH_2_O was added to the bottom of the tube for prehydration followed by 1 mL of 0.4 M sucrose. The solution was incubated for 30 minutes at 50 °C. After the solution cooled to room temperature (21-24 °C), the MmTMEM63B proteins were added to achieve the desired protein-to-lipid ratio, and the protein-lipid solution was shaken gently on an orbital mixer for an additional 3 hours at 4 °C. Giant artificial liposomes were observed in the recording bath following this incubation period. The protein-lipid solution (5-10 μL) was added to the bath buffer used for patch-clamp recordings and waited approximately 10 min for unilamellar blisters to form. For imaging of reconstituted liposomes, a protein-to-lipid ratio of 1:100 (wt:wt) was sufficient. Confocal images of liposomes were obtained using a confocal microscope (IXplore SpinSR, Olympus) with a 100× oil-immersion objective lens and an excitation wavelength of 488 nm.

### Patch-clamp recording

For the electrophysiological recordings in CHO cells, pCMV-TMEM63B-IRES-eGFP construct was transfected into CHO cells using lipofectamine 3000 (Thermo Fisher Scientific) according to the manufacturer’s instructions. Cells were seeded 24-48 h following transfection onto matrigel-coated coverslips and used for recordings 4-12 h. Whole-cell currents were recorded under a voltage clamp using 3∼5 MΩ borosilicate glass pipettes. Cells were continuously perfused (∼3 ml/min) at room temperature with isotonic extracellular solutions containing (mM) 140 NaCl, 5 KCl, 2 MgCl_2_·6H_2_O, 2 CaCl_2_·2H_2_O, 10 HEPES, and 10 D-glucose, pH 7.4, with NaOH (300 mOsm/Kg). Additional sucrose was added to increase the extracellular osmolarity without changing the ion concentration. The pipette solution contained (mM) 140 CsCl, 5 EGTA, and 10 HEPES. The osmolarities of all solutions were assessed by a freezing point osmometer (Gonotec, 3000 Basci, Germany). The whole currents were recorded with a Multi-Clamp 700B amplifier and Digidata 1550B digitizer processed by pClamp11.1 software (Molecular Device). Cells with a membrane resistance below 800 MΩ or series resistance above 10 MΩ were discarded. Currents were sampled at 10 kHz and low-pass filtered at 1 kHz. The cells were held at 0 mV before the application of test protocols. The protocol for measuring the hypertonic-activated currents was a 100 ms ramp from -80 to 80 mV every 2 s. Solution changes were performed manually using a 20 mL syringe. The whole-cell currents were recorded in the isotonic solution (300 mOsm/Kg) for 2 minutes and then switched to a hyperosmotic solution (320, 350, 400, 500 and 600 mOsm/Kg) for 3 minutes before isotonic wash-out.

For the electrophysiological recordings in SFO neurons, wild-type or *Tmem63b^HA-fl/HA-fl^* mice were injected with AAV-CaMK2α-mCherry or AAV-CaMK2α-Cre-mCherry virus in the SFO for at least two weeks. Mice were anesthetized with pentobarbital sodium (50 mg/kg) and then quickly decapitated to remove the brain into ice-cold cutting solution containing (in mM) 2 KCl, 1.25 Na_2_HPO_4_, 0.2 CaCl_2_, 12 MgSO_4_, 10 D-Glucose, 220 Sucrose and 26 NaHCO_3_ and oxygenated with 95% O_2_ + 5% CO_2_. Coronal brain slices including the SFO (300 μm) were prepared with a vibratome (VT1200S, Leica). After sectioning, the slices were incubated in standard artificial cerebrospinal fluid (ACSF) containing (in mM) 126 NaCl, 2.5 KCl, 1.25 Na_2_HPO_4_, 2 CaCl_2_, 10 D-Glucose, and 26 NaHCO_3_ were oxygenated with 95% O_2_ + 5% CO_2_ for 20-30 min at 34 °C. Then, the slices were allowed to recover for at least 1 h at room temperature before recording. Additional sucrose was added to increase the extracellular osmolarity without changing the ion concentration. Neurons exhibiting red fluorescence were chosen for recording. Action potentials were cell-attached recorded and when recording glutamate-evoked EPSC, cells were voltage-clamped at -70 mV in ACSF containing 10 μM SR95531 to block GABA_a_ receptors. Focal responses were evoked by exogenous glutamate applied toward the soma of SFO excitatory neurons via a patch pipette containing 50 μM glutamate with a pressure of 30 psi under the control of pressure application system (LHDA0533115H, The Lee Co., Westbrook, CT, USA). When recording hyperosmotic currents in SFO neurons, cells were voltage-clamped at -70 mV and was recording in ACSF containing 50 μM APV, 20 μM CNQX and 10 μM SR95531 to block glutamate and GABA_a_ receptors. Path-clamp recordings were performed using a Multi-Clamp 700B amplifier, with signals being recorded/analyzed using the Digidata 1550B data acquisition system and pClamp 11.2 software package (Molecular Devices).

For the electrophysiological recordings in the MmTMEM63B-reconstituted proteoliposomes, the protocol used to record currents from proteoliposomes was previously reported^45^. A 1:400 protein-to-lipid ratio (wt:wt) was used for current recordings. For the MmTMEM63B-eGFP channel recordings, the bath solutions contained (in mM) 140 KCl, 10 HEPES, 1 MgCl_2_, 10 glucose, pH 7.3 adjusted with KOH (300 mOsm/kg), and the internal solution contained (in mM) 130 NaCl, 5 KCl, 10 HEPES, 10 TEA-Cl, 1 CaCl_2_, 1 MgCl_2_, pH 7.3 adjusted with NaOH. For the hypertonic stimulus (320, 350, 400, 500 and 600 mOsm/kg) to obtain activated channel activity of MmTMEM63B, additional sucrose was added to the pipette buffer. For the blockage of channel activity, 30 µM GdCl_3_ was added to the pipette buffer. Borosilicate glass pipettes (BF150-86-10, Sutter Instrument) were pulled using a pipette puller (P-97, Sutter Instrument) to a diameter corresponding to the optimal pipette resistance, which was in the range of 4.0-6.0 M Ω . The patch resistance to proteoliposomes increased to more than 1 GΩ. Patch-clamp experiments in liposomes were performed in excised patch (inside-out for liposomes) mode using an EPC-10 amplifier and Patchmaster software (HEKA Electronic, Lambrecht, Germany), filtered at 2 kHz and digitized at 20 kHz. The channel recordings can be analyzed using pCLAMP 11.2 software, and data were plotted using GraphPad Prism 8.0. The current responses in the empty liposomes and MmTMEM63B-reconstituted proteoliposomes were counted under isotonic or hypertonic conditions holding at 40 mV. The integrated current was calculated as the area above baseline for a 10-second duration. All experiments were performed at room temperature.

### Statistical analysis

Statistical analysis was performed by IBM SPSS 22.0. Figures were plotted using GraphPad Prism 8.0. Corresponding statistical methods were selected according to different experimental designs and data types. An independent sample *t test* was used to compare two independent samples with linear parameters. *One-way* or *two-way* ANOVA was used for comparisons among multiple groups. The *LSD* method was used for pairwise comparison among multiple groups in the case of homogeneity of variances, while *Dunnett’s T3* method was used in the case of uneven variances. Data are presented as the mean ± SEM. Statistical significance is defined as \**p* < 0.05, \*\**p*<0.01, \*\*\**p*<0.001.

### Supplemental information

Figures S1-S11

## Supplemental Figure Legends

**Figure S1.**
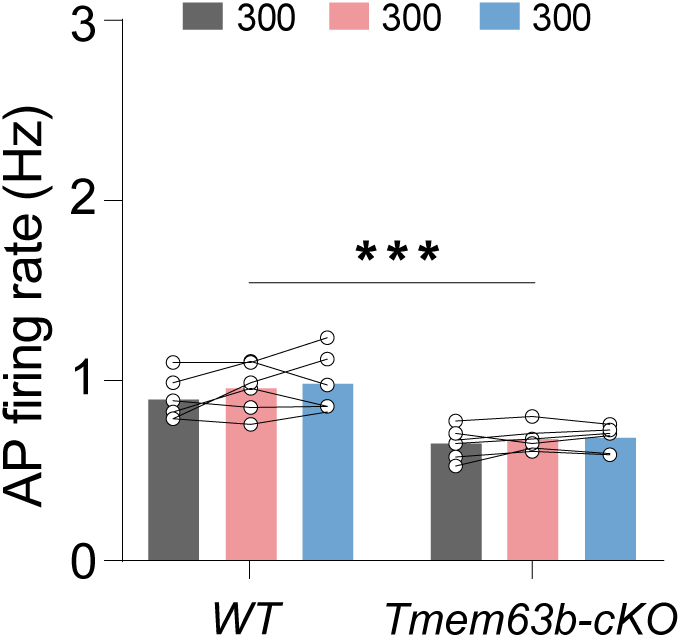
Action potential firing rate of SFO excitatory neurons under isotonic stimulus. The firing rate of SFO excitatory neurons of WT or Tmem63b-cKO mice show no significant change over time with isotonic solution perfusion (300 mOsm/kg).

**Figure S2.**
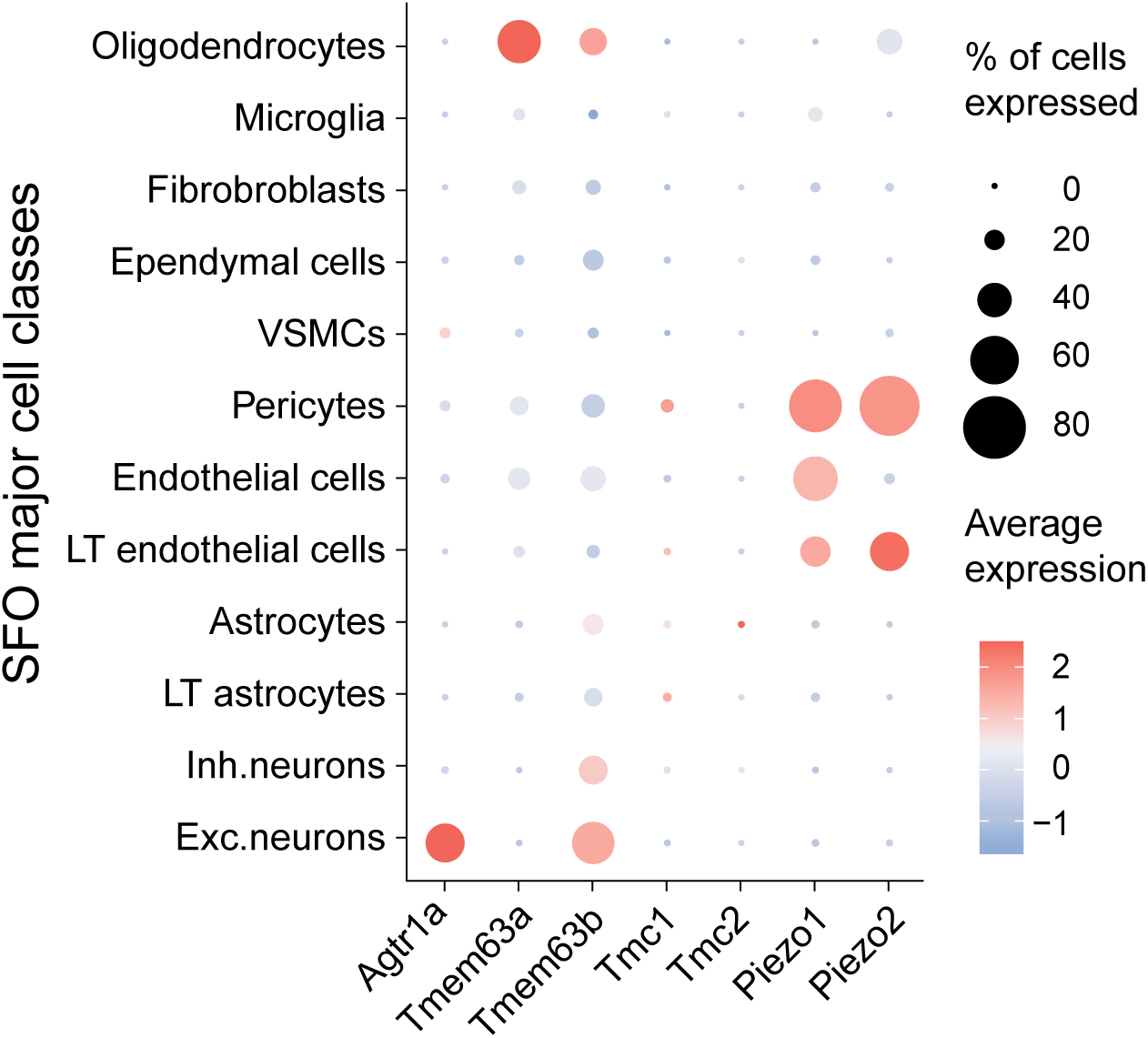
scRNA-Seq analysis of osmosensitive or mechanosensitive ion channel expression in the SFO. Dot plot of cell-type-specific expression for osmosensitive or mechanosensitive ion channels expression in the major cell classes of the SFO based on the published data17. Dot size is proportional to % of cells with transcript count >0 expression, color scale represents Z scored average gene expression, n=7950 cells.

**Figure S3.**
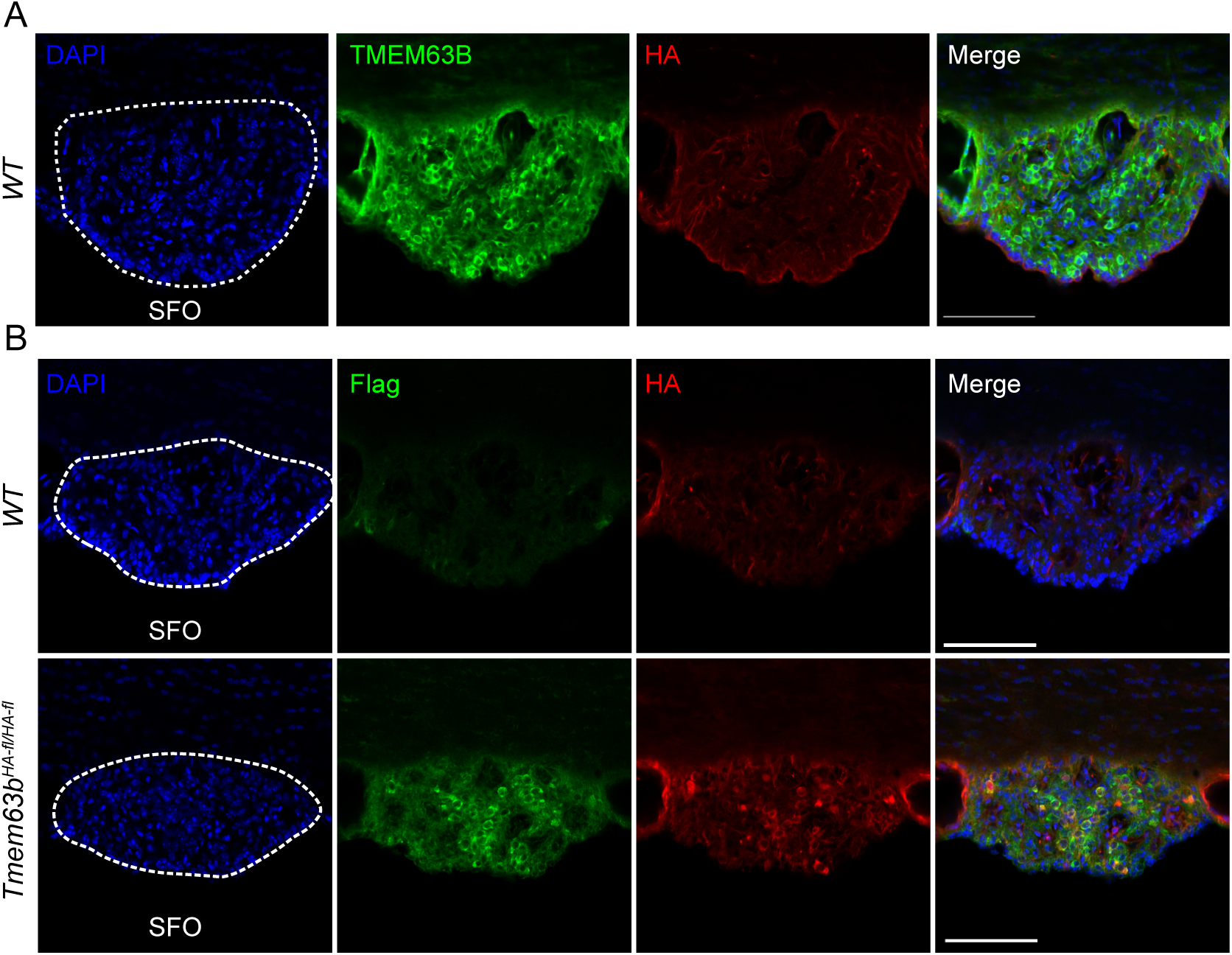
Protein expression pattern of TMEM63B in SFO. **(A)** Representative images show the expression of endogenous TMEM63B detected by anti-HA antibody and custom-made TMEM63B polyclonal antibody in the SFO of *Tmem63b^HA-fl/HA-fl^* mice. Scale bar 100 μm. **(B)** Representative images show the expression of endogenous TMEM63B in the SFO of *Tmem63b^HA-fl/HA-fl^* mice, stained with antibodies against a fusion HA (red) or 3×Flag (green) tag. WT mice show no signal in the SFO with these two antibodies (Lower panel). Scale bar 100 μm.

**Figure S4.**
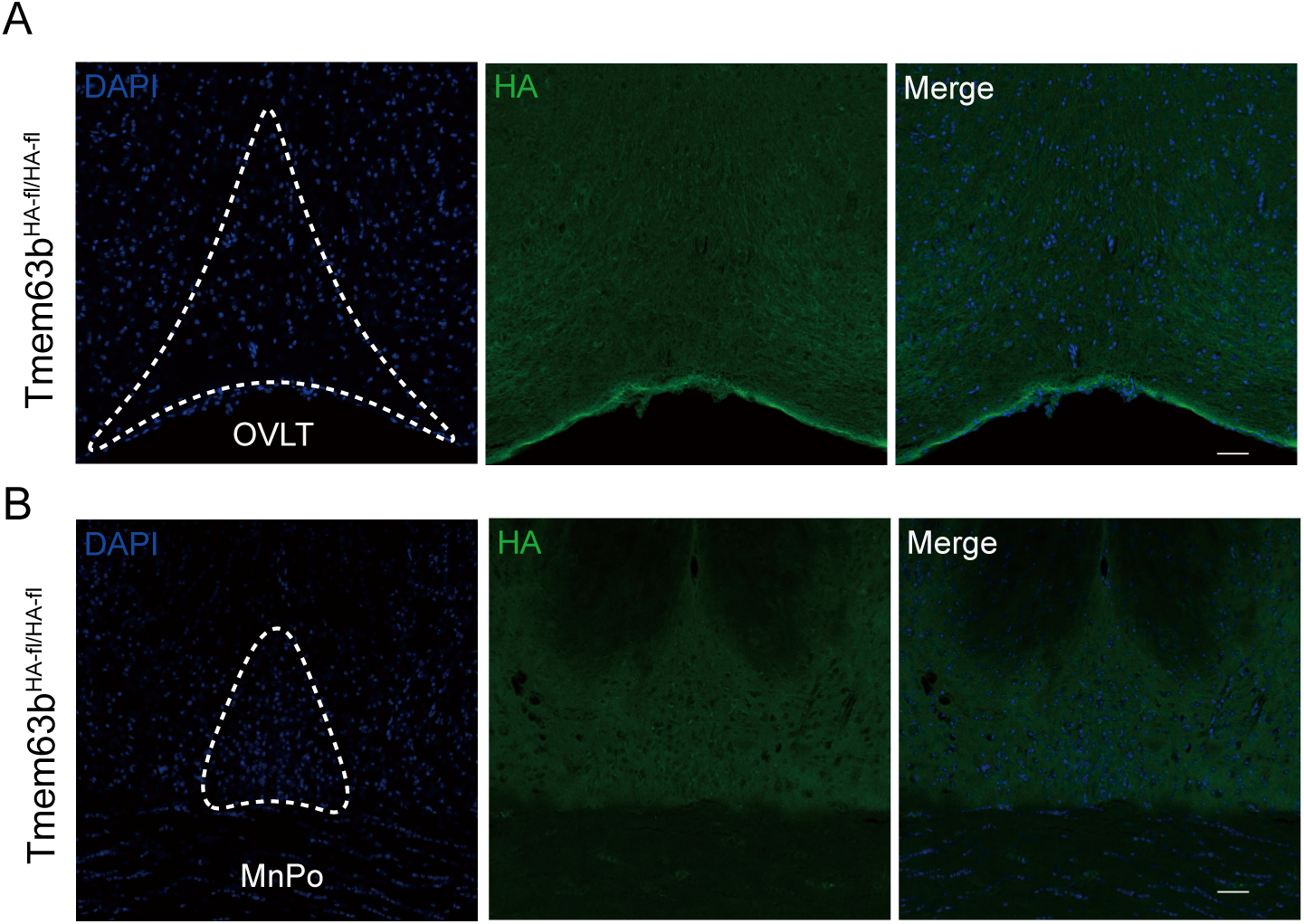
No endogenous expression of TMEM63B protein in OVLT and MnPO. **(A)** Representative images show no expression of endogenous TMEM63B in the OVLT of *Tmem63b^HA-fl/HA-fl^* mice, stained with antibody against a fusion HA (red). Scale bar 100 μm. **(B)** Representative images show no expression of endogenous TMEM63B in the MnPO of *Tmem63b^HA-fl/HA-fl^* mice. Scale bar 100 μm.

**Figure S5.**
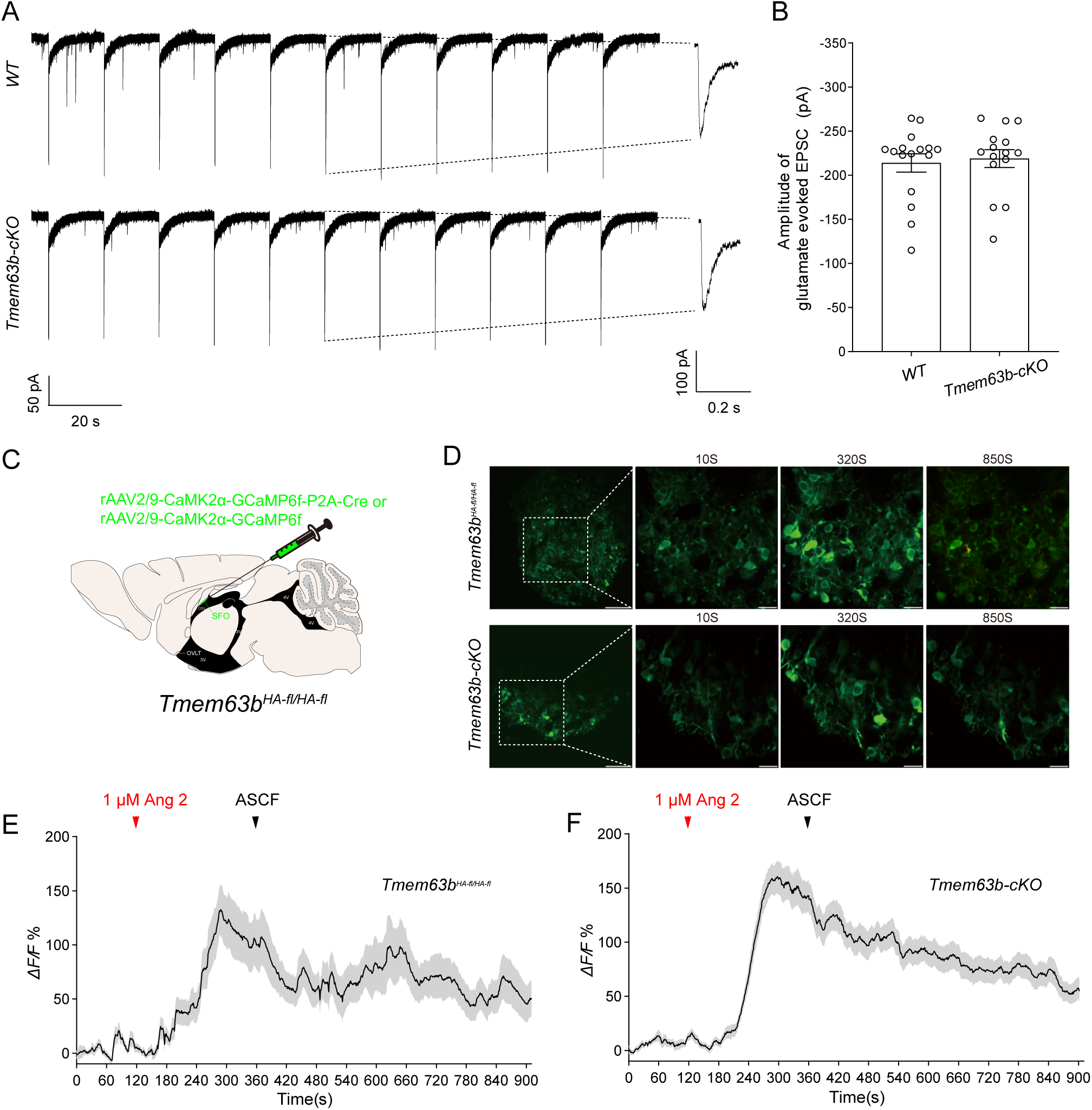
TMEM63B-deficient SFO excitatory neurons can respond to non-osmolarity stimulation. **(A)** Representative excitatory postsynaptic currents (EPSC) induced by puffing 50 µM glutamate 30 ms in every 15 s in the SFO excitatory neurons of *Tmem63b-cKO* mice. **(B)** Quantification of amplitude of glutamate evoked EPSC. All error bars denote the mean ± SEM, n =15-16. **(C)** Schematic diagram of injecting rAAV2/9-CaMK2α-GCaMP6f or rAAV2/9-CaMK2α-GCaMP6f-Cre in the SFO to expression GCaMP6f in SFO excitatory neurons in *Tmem63b^HA-fl/HA-fl^* mice. **(D)** Representative images showing SFO excitatory neurons calcium activity before, during and after bath application of 1 μM AngII in control and *Tmem63b-cKO* mice. **(E-F)** Similar calcium fluorescence intensity in SFO excitatory neurons in control and *Tmem63b-cKO* mice in response to 1 µM AngII application.

**Figure S6.**
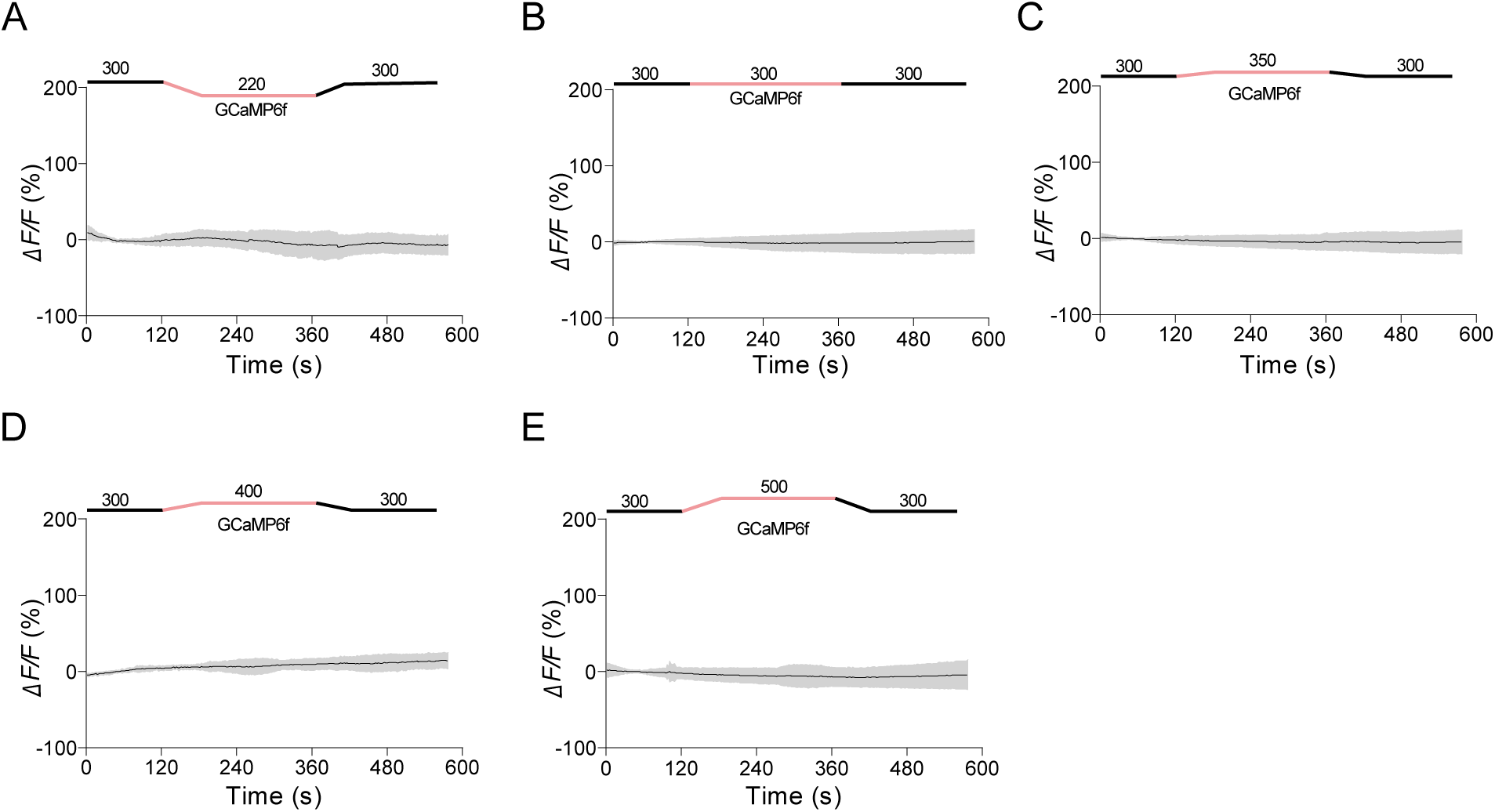
N2a cells transfected with GCaMP6f showed no significant calcium responses to osmolarity changes. **(A-E)** The calcium fluorescence intensity of N2a cells transfected with GCaMP6f, in response to hyposmolarity of 220 mOsm/kg **(A)**, isotonic solution **(B)** and hyperosmolarity of 350 mOsm/kg **(C)**, 400 mOsm/kg **(D)**, 500 mOsm/kg **(E)**. The red line indicates the hyperosmotic stimulus for 4 minutes. The grey shadow denotes the mean ± SD.

**Figure S7.**
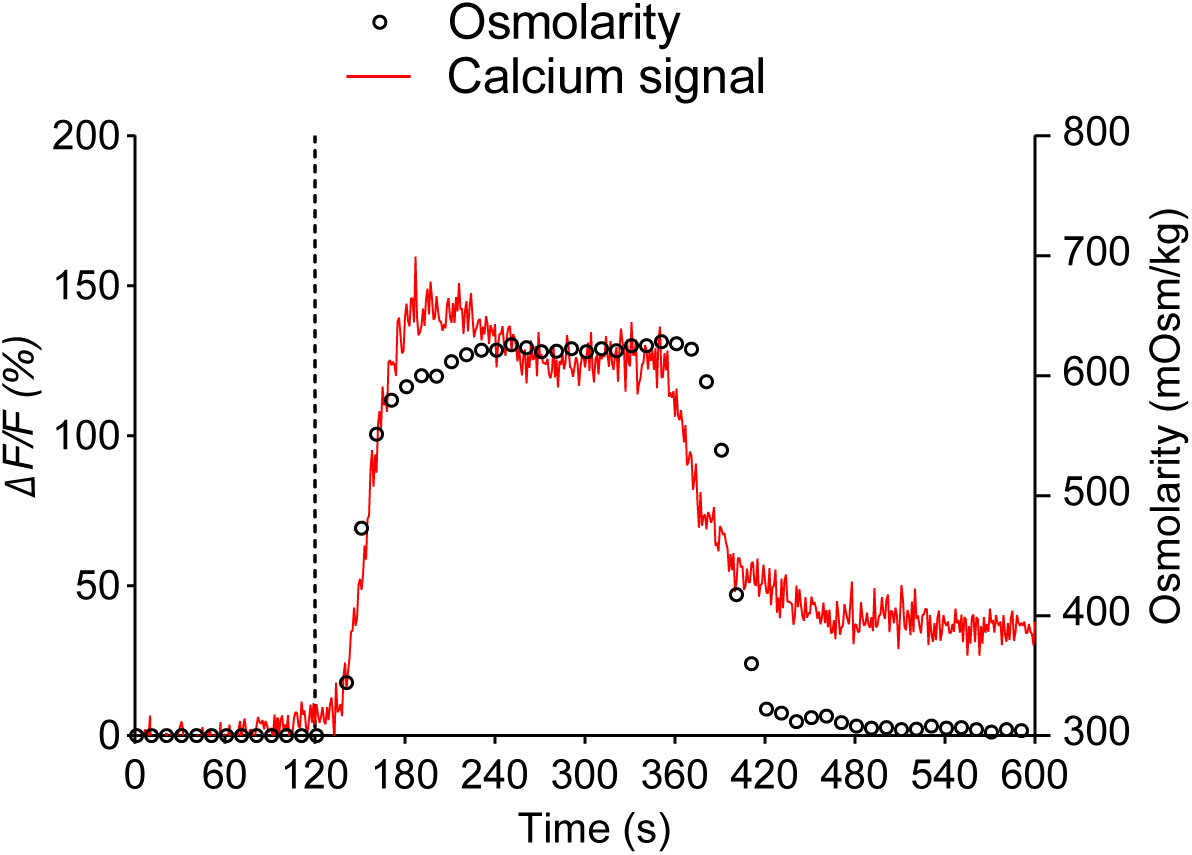
The calcium responses in TMEM63B transfected N2a cells coincides with the osmolarity changes. Calcium signal (red line) responses with osmolarity change (black dot) in an N2a cell expressing TMEM63B. The calcium signal was captured for 10 minutes at 0.5 Hz and the osmolarity of the external solution was measured every 10 seconds. The dashed line indicates the onset time of application of the hypertonic solution. The left y-axis shows the response of calcium signaling (*ΔF/F*), and the right y-axis represents the osmolarity of the external solution (mOsm/kg).

**Figure S8.**
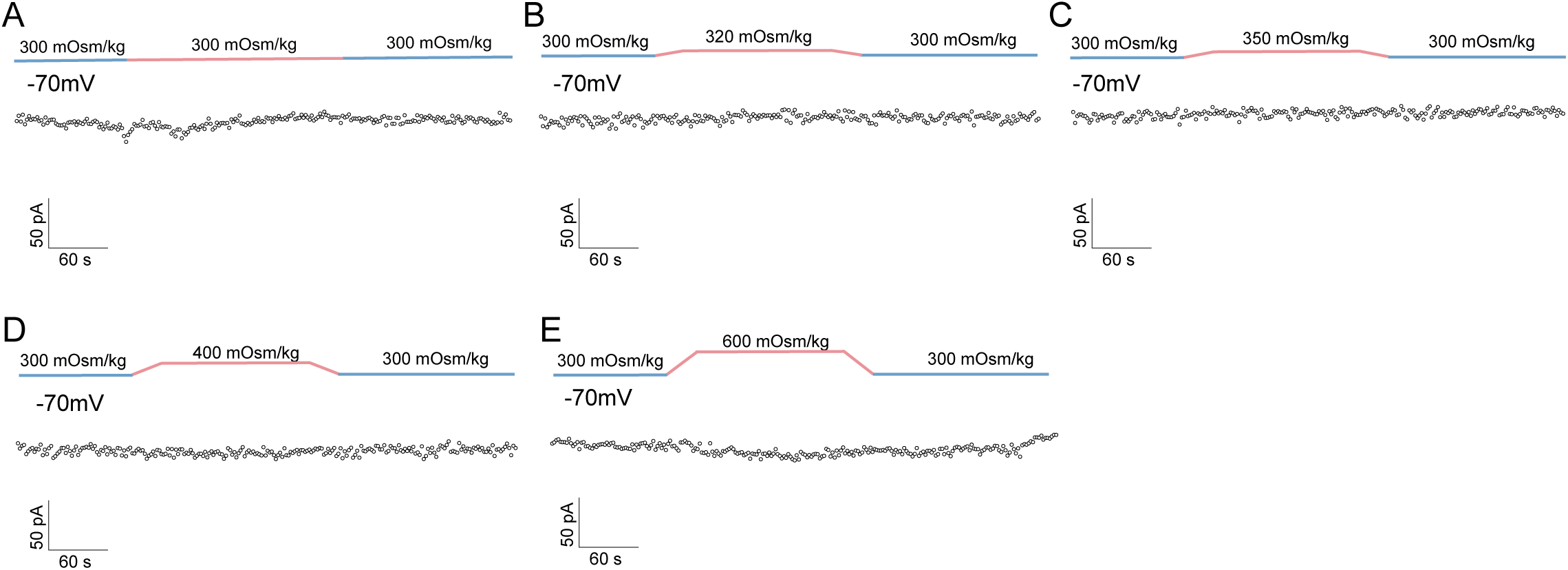
Untransfected CHO cells showed no significant hyperosmosensitive currents. **(A-E)** Representative whole-cell currents in untransfected CHO cells were recorded every two seconds by ramp protocols in 2-min isotonic solution (300 mOsm/kg; blue) followed by 3-min (red) of isotonic solution **(A)** or hypertonic solution of 320 mOsm/kg **(B)**, 350 mOsm/kg **(C)**, 400 mOsm/kg **(D)** and 600 mOsm/kg **(E)**, before isotonic wash-out. Currents at -70 mV over time were plotted.

**Figure S9.**
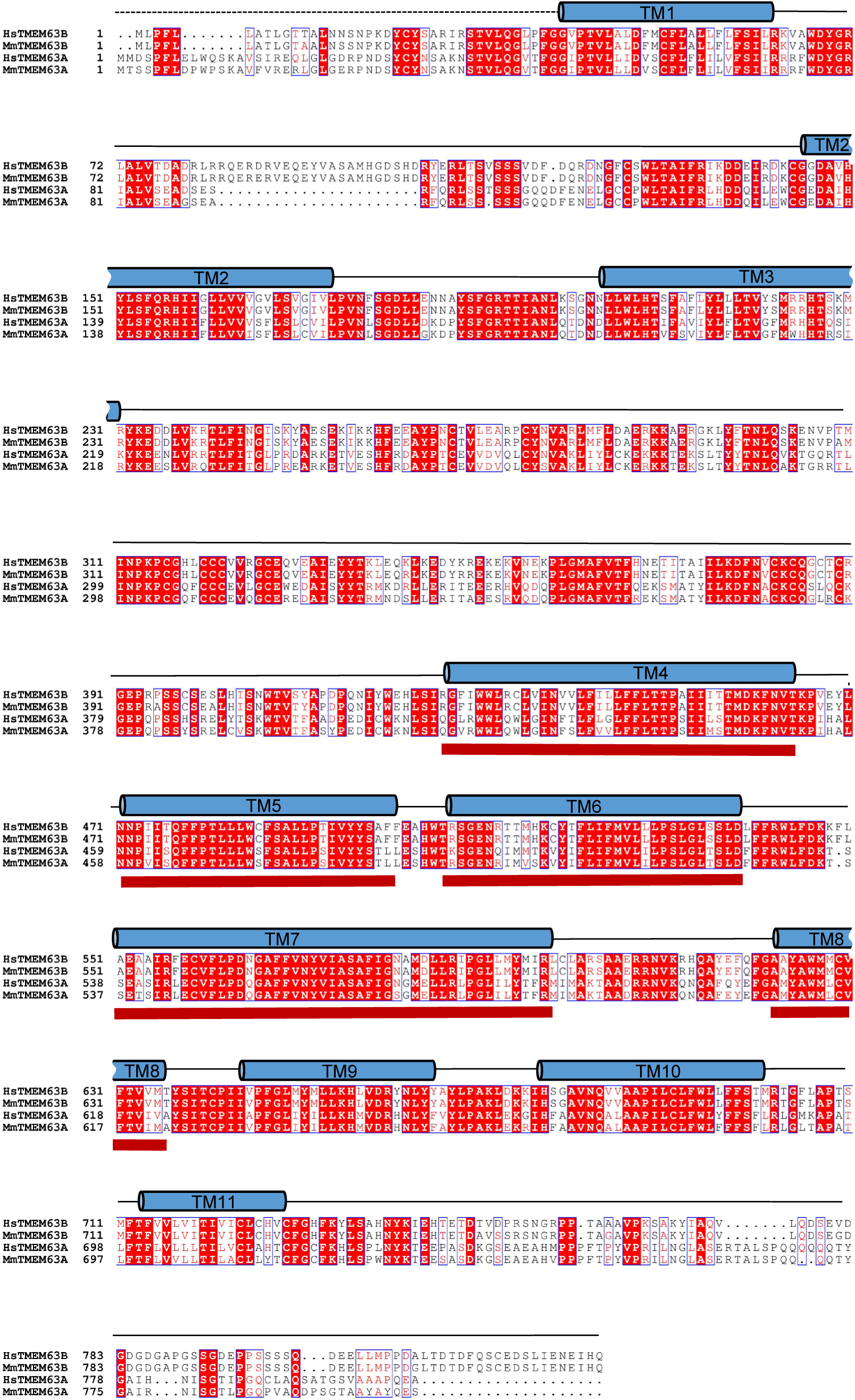
Alignment of amino acid sequences between human and mouse TMEM63 family proteins based on the model generated by SWISS-MODEL. The transmembrane regions is noted as blue cylinders, and identical residues are colored red, while similar residues are marked in red font. The red lines at the bottom of the sequences indicate the transmembrane helix around the putative pore region of HsTMEM63B, corresponding to the sequence of 426-460, 470-502, 506-539, 550-584, and 622-637.

**Figure S10.**
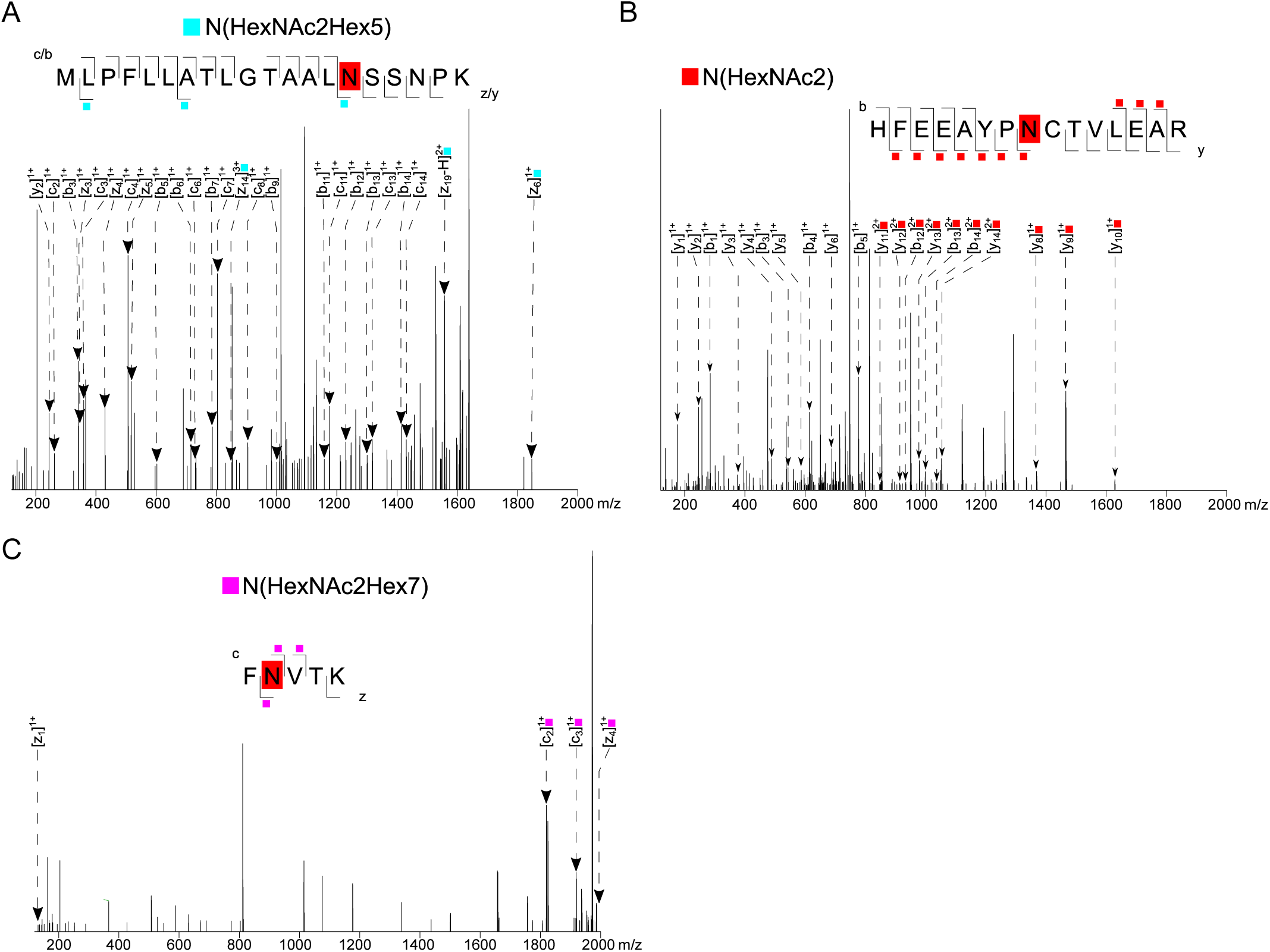
Mass spectrometry analysis of the glycosylation sites of MmTMEM63B. **(A)** The peptide sequence (MLPFLLATLGTAALNSSNPK) at position N15 featured the N-glycosylated oligosaccharide HexNAc2Hex5 (high-mannose N-linked glycan). **(B)** The peptide sequence (HFEEAYPNTVLEAR) at position N266 also presented the N-glycosylated oligosaccharide HexNAc2 (complex-type N-linked glycan). **(C)** The peptide sequence (FNVTK) at position N462 possessed the N-glycosylated oligosaccharide HexNAc2Hex7 (high-mannose N-linked glycan). By analysis of the mass spectrometry results based on the database, these glycan types were identified.

**Figure S11.**
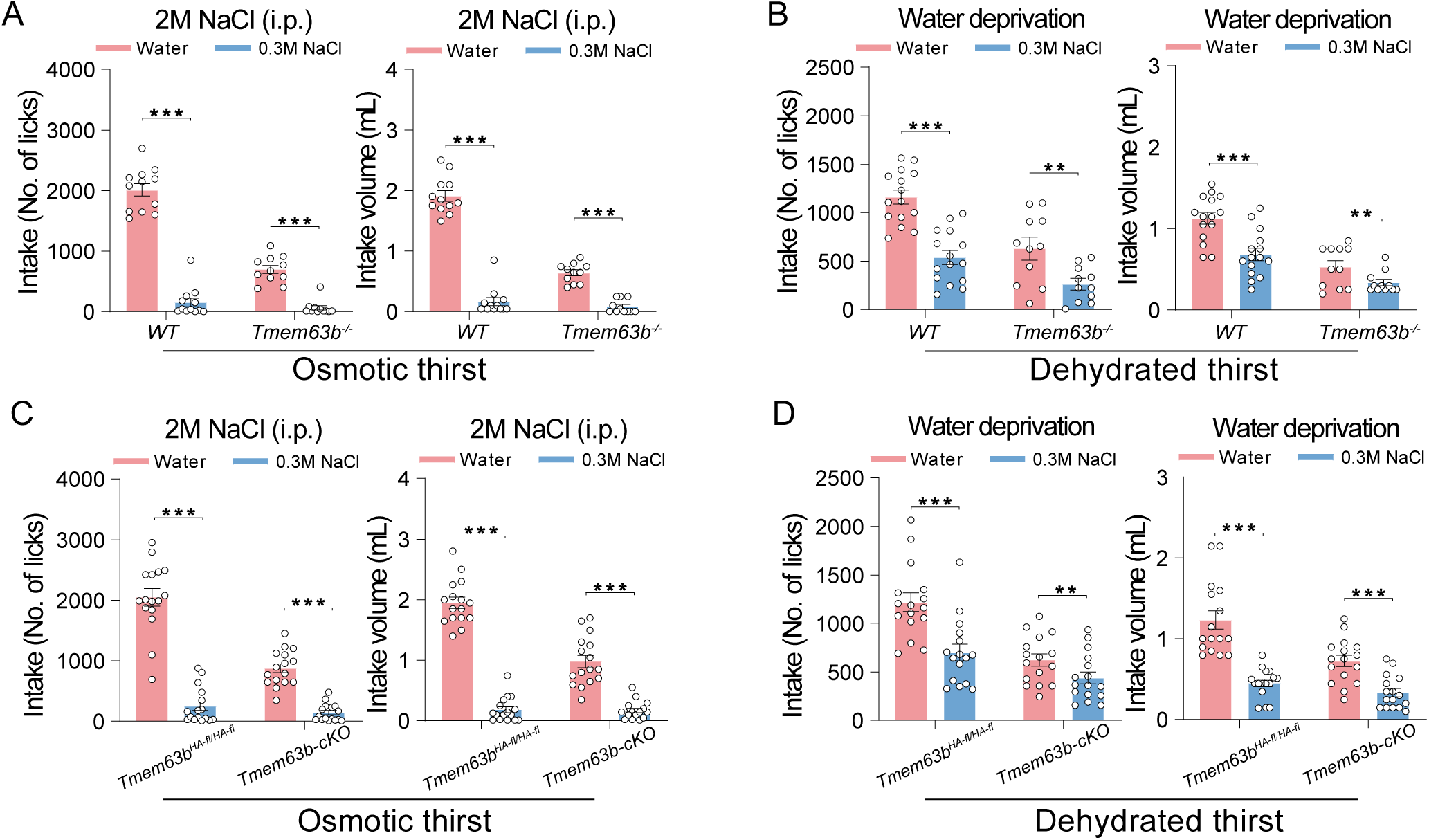
Water and hypertonic saline (0.3 M NaCl) intake of *Tmem63b-KO* and *Tmem63b-cKO* mice under osmotic and dehydrated thirst. **(A)** Water (red) and hypertonic saline intake (0.3 M NaCl, blue) of wild-type and *Tmem63b^-/-^* mice during a 1-h session after 2 M NaCl injection. Both WT and KO mice showed a strong preference for water, with little hypertonic saline intake under osmotic thirst. **(B)** Water and hypertonic saline intake of wild-type and *Tmem63b* knockout mice after water deprivation during a 1-h session. **(C)** Water and hypertonic saline intake of *Tmem63b^HA-fl/HA-fl^* mice and *Tmem63b^HA-fl/HA-fl^* mice injected with AAV-CaMK2α-Cre-mCherry virus under osmotic thirst during a 1-h session. Both *Tmem63b^HA-fl/HA-fl^* and *Tmem63b-cKO* mice show high water preference and barely consume 0.3 M NaCl. **(D)** Water and hypertonic saline intake of *Tmem63b^HA-fl/HA-fl^* mice and *Tmem63b^HA-fl/HA-fl^* mice injected with AAV-CaMK2α-Cre- mCherry virus under water deprivation during a 1-h session. In these figures above, water and 0.3 M NaCl intake are quantified by two parameters, number of licks (left) and water intake volume (right) during a 1-h session. All error bars denote the mean ± SEM, \*\*\**P*<0.001, Student’s unpaired *t* test, *n*=11-16.

